# Single-plant-omics reveals the cascade of transcriptional changes during the vegetative-to-reproductive transition

**DOI:** 10.1101/2023.09.11.557157

**Authors:** Ethan J. Redmond, James Ronald, Seth J. Davis, Daphne Ezer

## Abstract

Plants undergo rapid developmental transitions, as well as gradual developmental processes. Moreover, individual plants within a population will undergo the developmental transitions asynchronously, so it is difficult to assemble a time series to resolve the sequence of transcriptional changes that take place during these rapid transitions. Single-plant-omics has the potential to distinguish between transcriptional events that are associated with these binary and continuous processes. Furthermore, we can utilise single-plant-omics to exploit this developmental asynchrony to order individual plants by their developmental trajectory, revealing a detailed cascade of transcriptional events.

Here, we utilise single-plant-transcriptomics to resolve the transcriptional events that coincide with the onset of bolting. We performed RNA-seq on the leaves of individual plants from a large population of wild type Arabidopsis thaliana replicated at one time point during the vegetative-to-reproductive transition. Even though more than half of transcripts were differentially expressed between bolted and unbolted plants, we were able to find a subset of regulators that were more closely associated with gradual developmental traits like leaf size and biomass. Using a novel pseudotime inference algorithm, we determined that some senescence-associated processes, such as the reduction in ribosome biogenesis, are evident in the transcriptome before a bolt is visible.

These results show the potential of single-plant-omics to reveal the detailed sequence of events that occur during rapid developmental transitions.

**Graphical abstract:** Graphical Abstract: Physiological changes around bolting can be categorised into: ‘binary’ processes, which appear to have either occurred or not occurred at any given timepoint; or continuous processes, which can be observed quantitatively. For binary processes, expression of strongly correlated genes can appear to follow a ‘step’ change dynamic over time. However, when considered over a shorter timescale, the dynamics appear much smoother. For continuous processes, the shorter timescale should also capture smooth changes in gene expression.

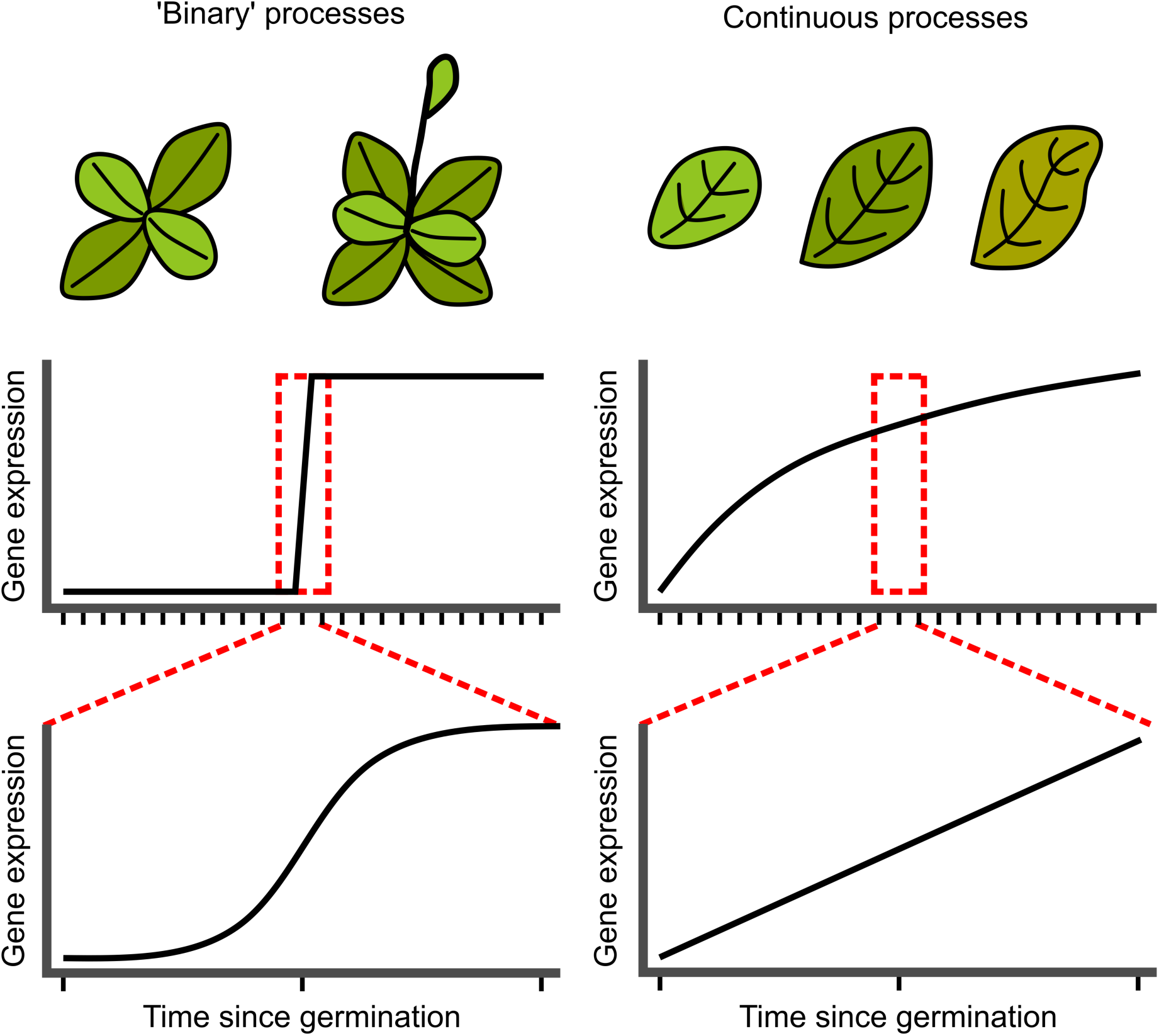

## Introduction

Plants experience many gradual developmental changes, such as leaf and biomass accumulation. However, at certain critical points, they undergo rapid ‘phase changes’ that act like switches, such as the juvenile-to-vegetative transition and the vegetative-to-reproductive transition (see Graphical Abstract). Transcriptional programmes coordinate these both gradual and binary developmental processes. Understanding the rapid transcriptional changes, such as those that occur during developmental transitions, requires a high temporal resolution of sample collection (Ezer and Keir 2019), a strategy that has effectively been deployed to detect rapid transcriptional response to the onset of light (Balcerowicz et al., 2021) and heat (Cortijo et al. 2017). However, this experimental approach does not easily translate to the study of late developmental transitions, because individual plants do not undergo developmental transitions in a synchronised way, and the degree of asynchrony is greater than the required temporal resolution of sampling. This developmental asynchrony – also known as ‘developmental instability’ (Klingenberg 2019) - is caused by the inherently stochastic nature of biochemical processes, such as transcriptional bursting (Leyes Porello, Trudeau, and Lim 2023). Developmental asynchrony is also affected by genetic factors and heterogeneity in microenvironments (Klingenberg 2019; Pertoldi et al. 2006).

In this manuscript, we suggest that single-plant-omics offers an innovative new way to investigate the transcriptional dynamics at developmental transition points. Single-plant-omics has previously been used to characterise diurnal gene expression noise (Cortijo et al. 2019) and maize transcriptional heterogeneity within a field-grown population (Cruz et al. 2020). By exploiting the developmental asynchrony across individual plants, we can reconstruct the order of transcriptional cascades that occur during a rapid developmental transition.

We focus on the vegetative-to-reproductive transition, an irreversible phase change in angiosperms that includes the formation of a bolt and eventually leads to flowering. Climate change has already been shown to reduce the synchrony of flowering time, leading to ecological and agricultural repercussions (Zohner, Mo, and Renner 2018). To support the initiation of reproductive growth, multiple processes are tightly coordinated within leaves during this transition. Of particular importance is the onset of leaf senescence: a transcriptionally-regulated and tightly controlled process that leads to cell death in leaves (Kim, Kim, et al. 2018) and nutrient reallocation to reproductive structures (Havé et al. 2017). Bolting-associated leaf senescence is an important contributor to the nutrient quality of grains (Havé et al. 2017; Gregersen, Holm, and Krupinska 2008).

In this paper, we utilise single-plant-omics of a population of wild type plants to reconstruct the transcriptional cascades that control development during the onset of bolting in Arabidopsis leaves. We find that the majority of genes are differentially expressed during bolting, reflecting a binary shift in transcriptional states. Next, we identify a subset of genes that are more closely associated with the gradual transcriptional changes associated with growth. Finally, we order the plants on the basis of their biological age, which enables us to uncover the sequence of events that occur during the bolting process, which begins with a shutdown of ribosome production and ends with the shutdown of photosynthesis. The gene network that we infer can serve as a reference for understanding the transcriptional regulation pathways that underlie this critical developmental transition and highlight the power of single-plant-omics in understanding complex developmental phase transitions.

## Results

### 1: Widespread transcriptional changes occur after bolting in *Arabidopsis thaliana*

Bolting functions as an indicator of the vegetative-to-reproductive developmental transition and occurs simultaneously with the onset of leaf senescence and the cessation of leaf development (Hinckley and Brusslan 2020; Möller-Steinbach, Alexandre, and Hennig 2010). To investigate the gene expression changes that arise during bolting, we measured gene expression from leaves of individual plants within a large population of wild type *Arabidopsis thaliana,* grown under uniform, controlled conditions (see Methods), sampled when a substantial proportion of the plants had bolted. To explore the connection between asynchronous gene expression and the major developmental changes occurring around the vegetative-to-reproductive transition, we recorded key developmental physiological traits from each plant.

We observed that plants with the same bolting status have more closely correlated transcriptomes than plants with differing bolting statuses (Supplementary Figure 2). We thus sought to characterise the transcriptomic changes occurring simultaneously with bolting, by finding differentially expressed genes (DEGs) between non-bolted and bolted plants, taking into account the large number of samples (see Methods and Table S1). Overall, 3754 genes were up-regulated and 6810 genes were down-regulated after bolting, accounting for 54% of the entire transcriptome after initial filtering (Supplementary Figure 1).

We performed GO term enrichment analysis on the DEGs (Raudvere et al. 2019). Significantly enriched terms cover a range of senescence-associated processes, consistent with previous observations that transcriptional changes during bolting are associated with senescence (Table S2) (Hinckley and Brusslan 2020).

Firstly, programmed cell death (GO:0012501) is overrepresented among genes that were up-regulated in bolted plants (Figure 1A (i)). Conversely, regulation of cell cycle (GO:0051726) is overrepresented within down-regulated genes (Figure 1A (ii)). These results are consistent with changes in cell cycles within maturing leaves. A shift occurs from cell division to cell elongation (marked by endoreduplication) in young to mature leaves, and eventually programmed cell death occurs in senescent tissue (Del Prete et al. 2019; Woo et al. 2019). Secondly, bolting correlates with a reduction of gene expression for genes associated with each step of transcription and translation, including RNA polymerase complex formation (GO:0030880), RNA processing (GO:0006396), and structural constituents of the ribosome (GO:0003735). In contrast, genes related to post-transcriptional modification, such as ubiquitination (GO:0016567) and protein phosphorylation (GO:0006468), were more expressed in bolted plants. Thirdly, photosynthesis-related (GO:0015979) gene expression was reduced in bolted plants (Figure 1A (iii)), consistent with chlorophyll catabolism during senescence (Woo et al. 2019). Finally, we observed an overrepresentation of genes associated with jasmonic-acid (GO:0009753), abscisic acid (GO:0009737) and salicylic acid (GO:0009751). These hormone signalling pathways were previously associated with leaf senescence (Woo et al. 2019). These results indicate that the onset of bolting coincides with a large-scale transcriptomic shift towards senescence.

**Figure 1:**
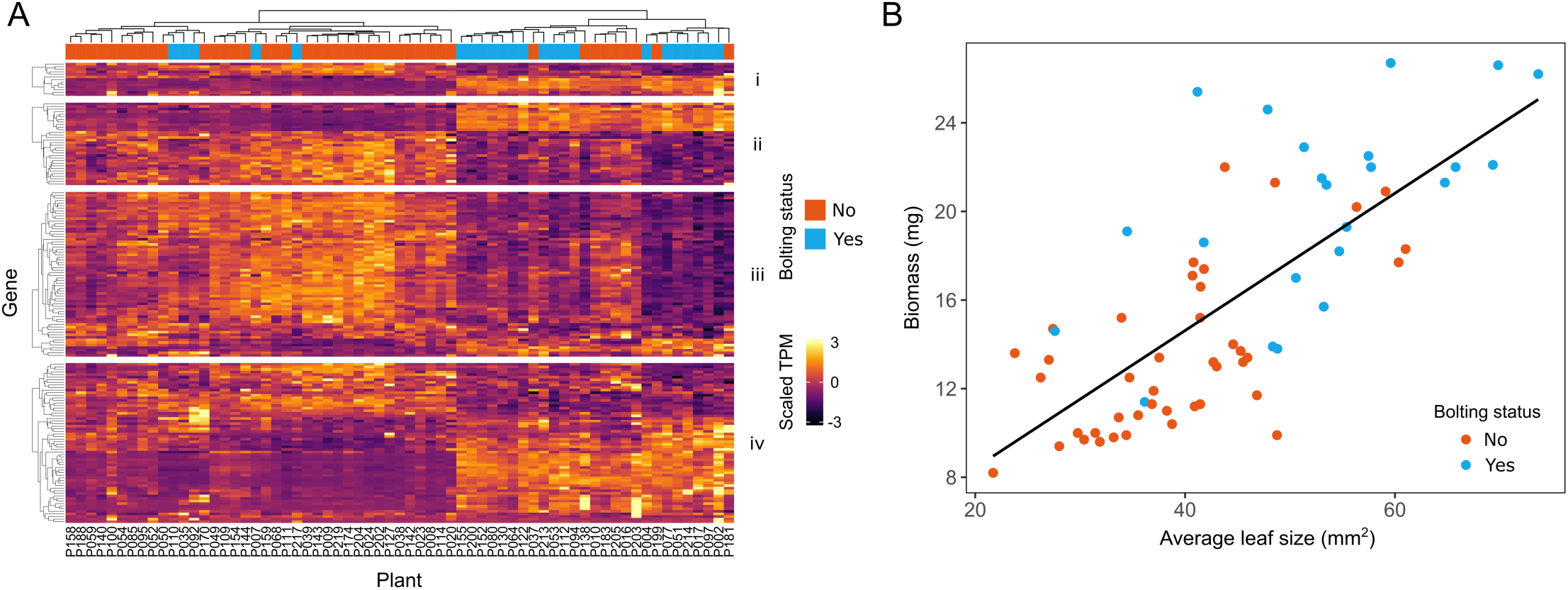
**A**: Gene expression values across all samples for genes annotated with selected GO terms. Only genes which were identified as differentially expressed are shown. Hierarchical clustering across each sample shows that plants with the same bolting status often cluster together. Additionally, hierarchical clustering within each group of genes indicates the nature of the large changes between non-bolted and bolted plants. i: Programmed cell death (GO:0012501). Adjusted P values: up = 2.68e-9, down = 1. ii: Regulation of cell cycle (GO:0051726). Adjusted P values: up = 1, down = 8.82e-15. iii: Photosynthesis (GO:0015979). Adjusted P values: up = 1, down = 3.84e-8. iv: Response to jasmonic-acid (GO:0009753). Adjusted P values: up = 1.97e-8, down = 1. Note that we scaled TPM values (to have a mean of 0 and standard deviation of 1 across all samples) then clipped any value above 3 or below -3. Adjusted P-values were calculated by g:Profiler and relate to test of GO term over-representation in either up- or down-regulated genes (Kolberg et al. 2020). For all hierarchical clustering, we used the ‘complete’ linkage method. **B:** Biomass and leaf size are correlated across all samples. Both leaf size and biomass are larger for bolted plants rather than non-bolted plants on average. Linear regression: R&#x005E;2 = 0.55, p = 1.59e-12.

### 2: Physiological traits have distinct correlations with biological processes

Bolting is a rapid developmental transition that occurs over a two- or three-day period, but other plant traits accumulate gradually over a plant’s lifetime, including leaf area and biomass. We observed that biomass and leaf size are correlated across the population, with larger plants more likely to have bolted (Figure 1B)

While we previously observed widespread transcriptional changes associated with the bolting transition, we hypothesised that there would be separate transcriptional programmes associated with leaf area and biomass. To investigate this, we used elastic net models to predict physiology from gene expression for all the plants in our population. We used leave-one-out-cross-validation (LOOCV) to validate the generalisability of our models. The performance of our prediction, as judged by the R&#x005E;2 scores between predicted and true measured values, was similar to another single-plant-omics study in maize (Figures 2A and 2B) (Cruz et al. 2020). We did not see a meaningful reduction of performance when reducing the set of predictors to only regulatory genes (see Methods and Supplementary Figure 4). Therefore, we only considered regulatory genes in the remainder of this section.

**Figure 2:**
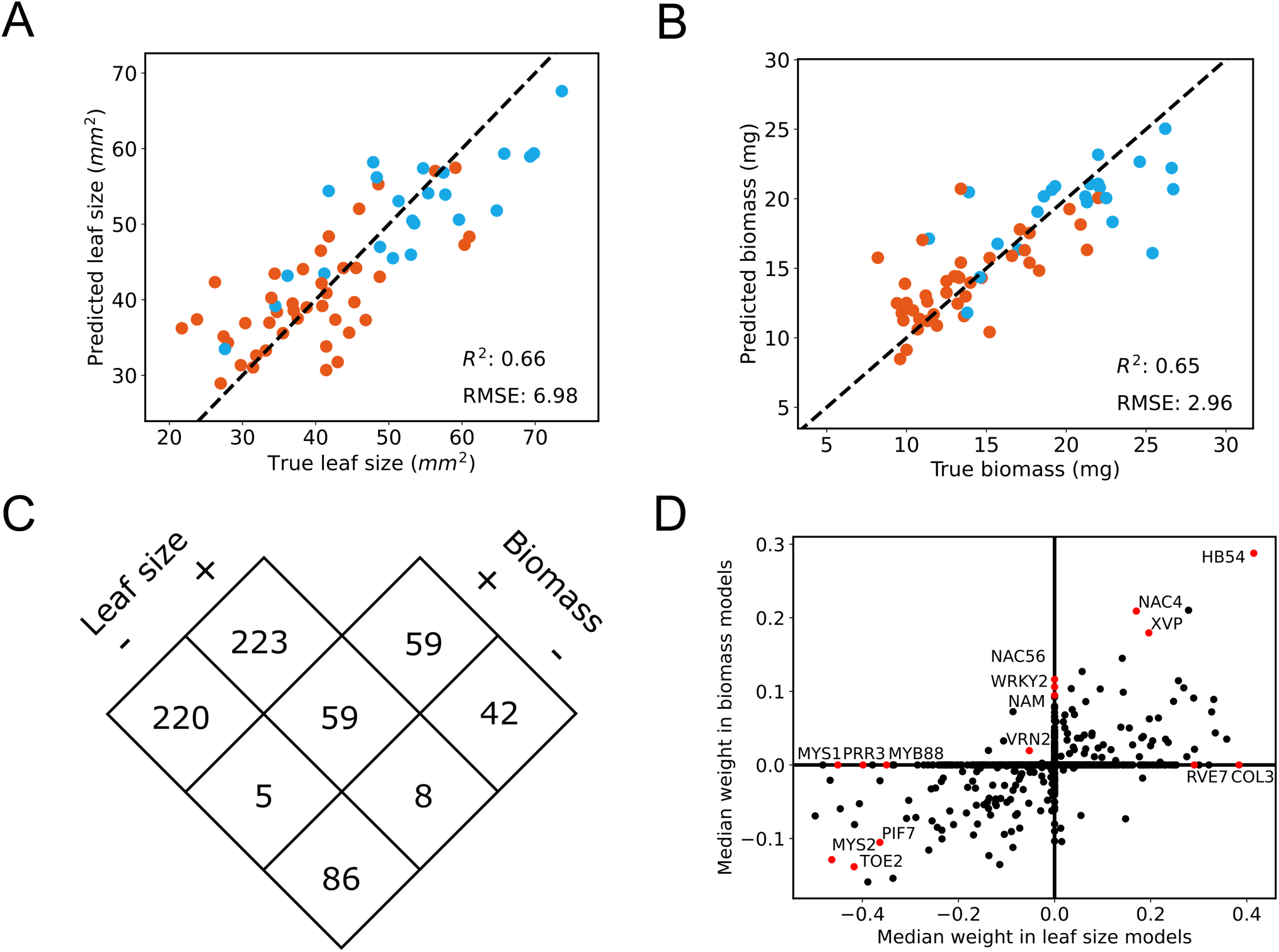
Figures about physiology prediction from gene expression. For **A** and **B**, red represents non-bolted plants and blue represents bolted plants. **A:** Performance of leaf size prediction from regulatory genes. Predicted values were produced from an Elastic Net model, trained on all data except for the relevant sample (see ‘Physiology trait prediction with Elastic Nets’ in Methods for more information on cross-validation). **B:** A repeat of A for biomass values. **C:** Overlaps between positive and negative predictors for leaf size and biomass Elastic Net models. Predictors were only included if they had a non-zero weight in every model during LOOCV cross-validation. **D:** Comparison of the median weight of predictors from biomass and leaf size models.

We were able to identify genes that were consistently identified as predictors of both leaf area and biomass in multiple models during the LOOCV process. Overall, 702 regulatory genes had a non-zero median coefficient across all models (Figure 2C). Of these, 59 predictors were identified as positively associated with biomass and leaf size – i.e. positive predictors – and 86 were identified as negatively associated with both traits, i.e. negative predictors (Figure 2C). The gene with the highest positive coefficient for both traits was *HOMEOBOX PROTEIN 54 (HB54;* AT1G27045), involved in a nitrogen-signalling cascade linked to plant growth (Ariga et al. 2022). Other genes highlighted in Fig 2C and discussed in the Supplementary Materials, suggest that these gradual traits are associated with transcription factors involved in the circadian clock, photomorphogenesis, wax biosynthesis and the microRNA/SPL-mediated juvenile to mature transition.

These results suggest that a small subset of the genes that are differentially expressed between bolted and unbolted plants are predictive of the continuous traits that are associated with plant growth.

### 3: Single-plant-omics can be used to generate age-related timeseries

The above results suggest that some genes are associated with continuous changes in the plant like biomass and leaf area, while others are associated with the ‘binary’ shift in gene expression from non-bolted to bolted plants. However, we expect that even a rapid developmental transition will involve continuous changes in gene expression near the transition point, but that observing these changes would require sampling many time points during the rapid transition period (Ezer and Keir 2019).

Although the individual plants in our population are the same chronological age, we observed that they are developing asynchronously, which we infer will lead to a diversity in the biological ages of the plants. We hypothesised that individuals within a population of plants would demonstrate asynchronous gene expression dynamics (Figure 3A), which would allow us to order the plants by their biological age. Here, we introduce a new method to order the plants by their biological age, based solely on gene expression data.

**Figure 3:**
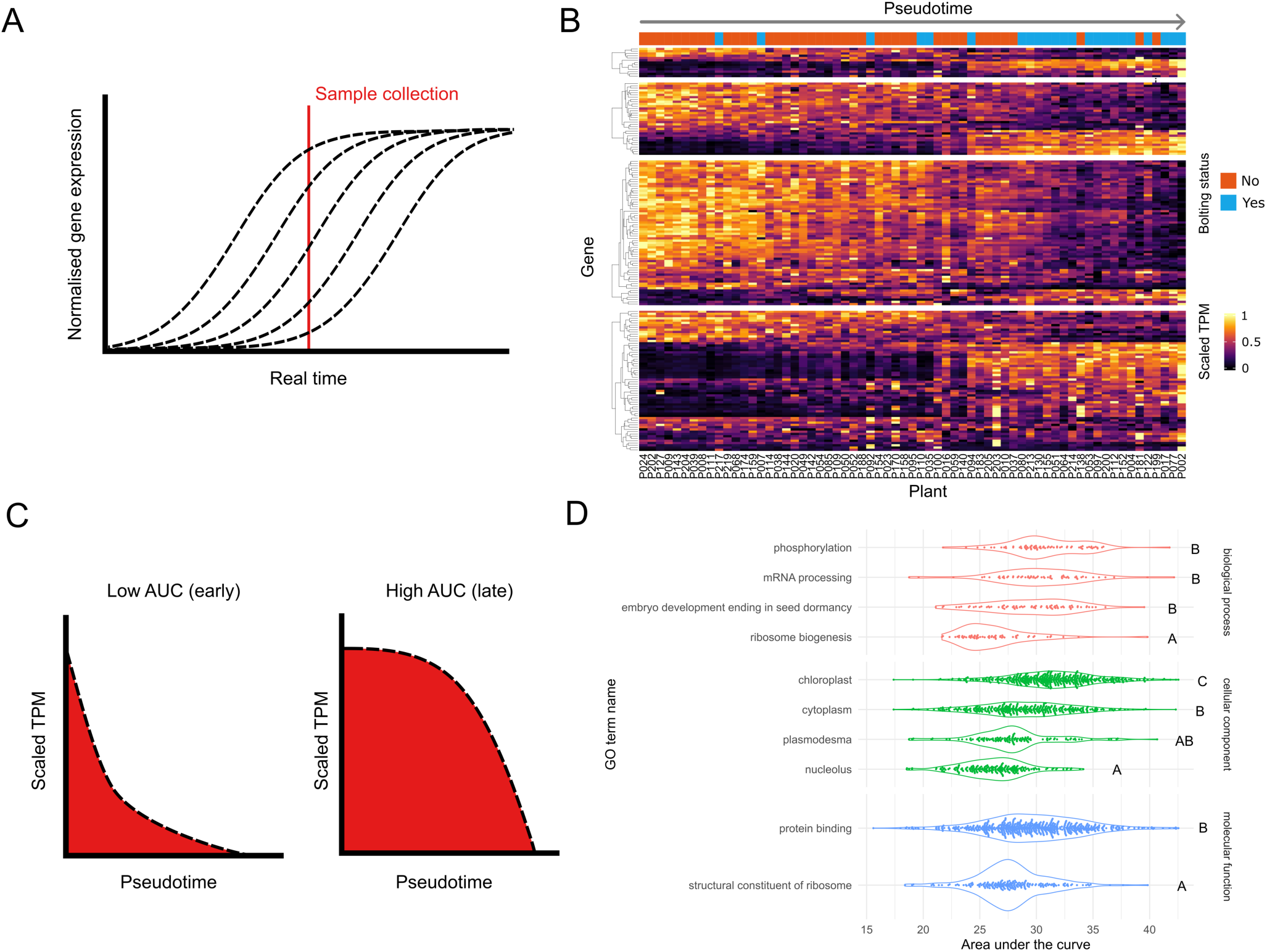
**A:** A theoretical interpretation of our sampling method. The dashed curves represent gene expression from individual plants across real time since germination. The red line represents the time when all samples were collected. Due to the desynchronisation of gene expression between plants, the red line captures a variety of gene expression values across samples. **B:** A repeat of the heatmap from Figure 1, reordered over pseudotime and with a different scaling of TPM values (see “Pseudotime Inference” in Methods). Gene expression dynamics are mostly monotonic and smoothly-changing. **C:** A diagram which demonstrates the area under the curve (AUC) values, for genes with decreasing gene expression. Dashed lines represent the theoretical gene expression values (after they have been scaled). The red region measures AUC. Genes whose expression decreases early on in pseudotime will have a low AUC value. Conversely, genes which decrease later on in pseudotime will have a high AUC value. **D:** AUC plots of key GO terms. These have been grouped by GO category (biological process, cellular component, and molecular function). Each dot represents a single gene annotated with that GO term. Letters depict significantly different GO terms based on

This problem of ordering samples based on gene expression data alone is related to *pseudotime inference* in the analysis of single-cell RNA-seq data. In that context, ‘pseudotime’ refers to an pseudotemporal ordering of single cells, which is assumed to contain information about a biologically meaningful developmental trajectories, such as cell differentiation (Saelens et al. 2019). Although multiple methods exist for pseudotime inference, the most popular approaches require large numbers of samples (cells) due to their unsupervised approach (Saelens et al. 2019). Notably, our sample size of 65 plants is lower than the (tens or hundreds of) thousands of cells required for unsupervised methods. To circumvent this restriction, we incorporated an assumption that the expression of most genes across the vegetative-to-reproductive transition is either monotonically increasing or decreasing. Our basis for making this assumption is our observation in Section 1 that the majority of genes are differentially expressed during bolting, implying that they primarily increase or decrease in expression. We were able to consistently order our plants along pseudotime to maximise the monotonicity of gene expression, using different subsets of genes (bootstrapping). Finally, we compiled a consensus pseudotime of samples after bootstrapping (Methods; Supplementary Figure 5).

We observed that pseudotime captures the continuous changes in gene expression of bolting-related processes, demonstrating the transcriptional shift during bolting is not a binary process. We visualised genes associated with programmed cell death, regulation of cell cycle, photosynthesis, and response to jasmonic acid over pseudotime (Figure 3B). Within each of these groups, there were clusters of genes with different expression timings, with changes in expression happening early or later in the bolting process. We defined a metric to describe whether genes were early or late changing: “area under the curve” (AUC; Figure 3C; Methods). For further analysis, we selected the top genes which varied smoothly over pseudotime (see Methods and Supplementary Figure 7 for details).

To better characterise the timing of different biological processes across the bolting transition, we tested for GO terms with significant differences between AUC values (Figure 3D; Supplementary Figures 9 and 10; Table S8; Methods). To understand the order of processes shutting down during the onset of senescence, we focused on genes that decreased their expression over pseudotime. Firstly, we found that ribosome biogenesis decreases earlier than processes related to phosphorylation, mRNA processing and embryo development leading to seed dormancy, suggesting that ribosomal production is slowed even before the onset of a visible bolt (Figure 3D). Similarly, terms for different cellular components decrease in the following order: nucleolus, cytoplasm, then chloroplast. Interestingly, the term plasmodesma decreases significantly earlier than chloroplast, suggesting that intercellular processes shut down before photosynthesis-related processes. Finally, we observe that mRNA binding decreases sooner than protein binding. These results suggest that ribosome production is one of the first processes to shut down at the onset of senescence, while photosynthesis is one of the last processes.

### 4: Pseudotime assists in the inference of gene regulatory networks

Our next aim was to identify the regulatory cascade that led to the large-scale transcriptional changes during bolting. We inferred gene regulatory networks (GRNs) from the gene expression data over pseudotime (Figure 4A, Table S10). To enrich for direct regulatory interactions, we filtered for potential regulatory links which were validated by DAP-seq (Methods; (O’Malley et al. 2016)). We only included transcription factors in our final network whose expression changed smoothly over pseudotime (see “Pseudotime Inference” in Methods) and that had DAP-seq data available for validation. The transcription factors in our network displayed varied expression patterns over pseudotime (Figures 4B and 4C), and they are ordered by their AUC in the network in Figure 4A. We identified four members of the SQUAMOSA PROMOTER BINDING PROTEIN-LIKE (SPL) family as potential regulators in our GRN. Two of these (SPL9, AT2G42200, and SPL13B, AT5G50670) are known to be targeted by miR156 during the vegetative-to-reproductive transition (Xu et al. 2016). Further, members of the NAC and WRKY transcription factor families are present among the putative regulators (Olsen et al. 2005; Phukan, Jeena, and Shukla 2016).

**Figure 4:**
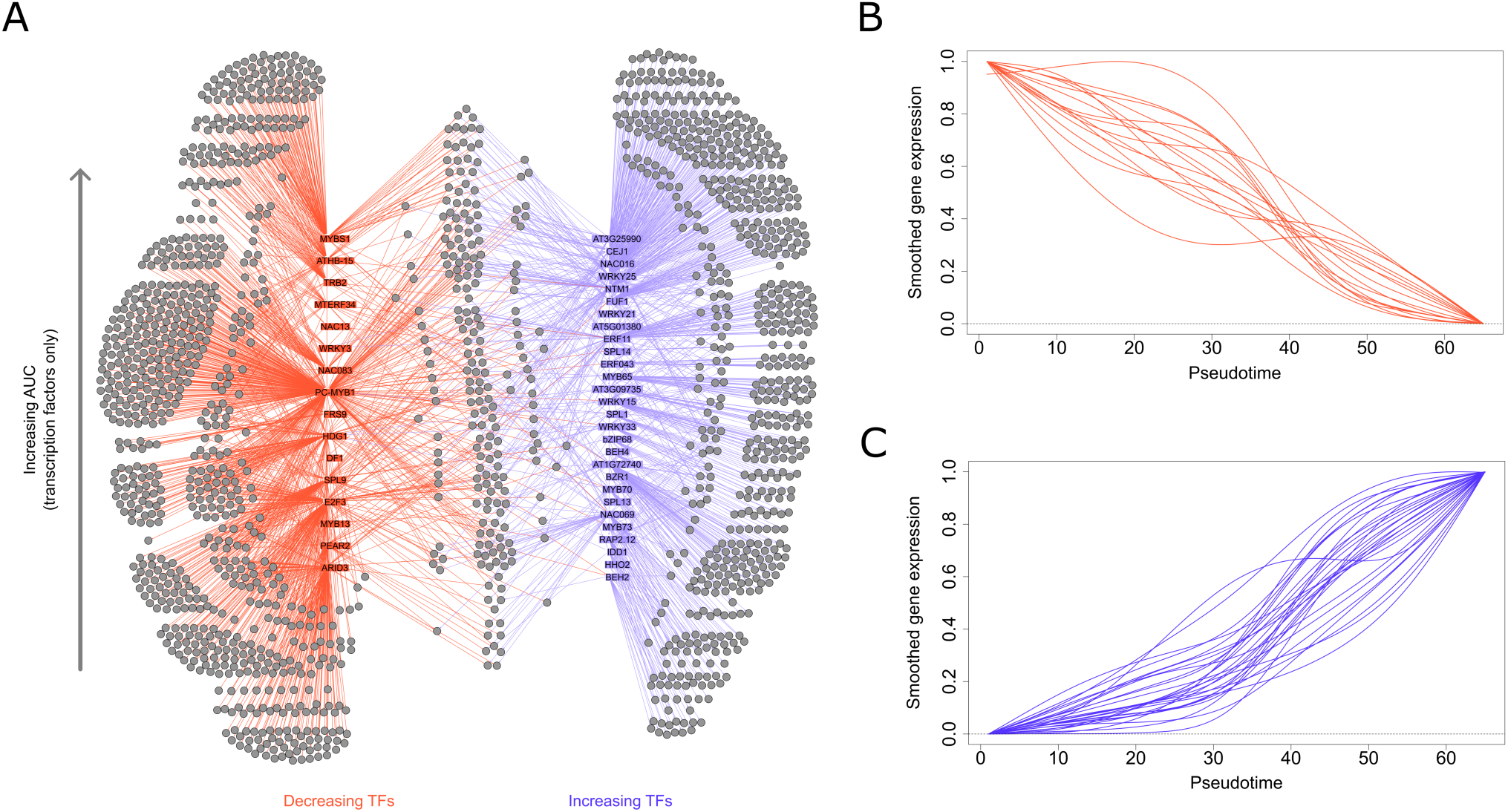
**A:** A visualisation of the gene regulatory network predicted by DynGENIE3, after filtering with DAP-seq data (see main text). Decreasing regulatory transcription factors are coloured in red, increasing ones in blue. Additionally, transcription factors are ordered bottom-to-top by increasing ranked AUC value. **B:** The smoothed gene expression values over pseudotime of decreasing TFs from the GRN. **C:** The smoothed gene expression values over pseudotime of increasing TFs from the GRN.

### 5: A small number of genetic variants are associated with pseudotime

Developmental asynchrony can be caused by environmental and genetic factors, as well as random stochasticity. Even in a nearly isogenic wild type population, there will be a number of genetic variants. In order to evaluate the role of genetics in encoding developmental asynchrony, we sought to determine the extent to which these genetic variants correlated with physiological traits and pseudotime. Within our population, we identified a total of 998 high-confidence variants (Figure 5A, see Methods for details on Variant Calling). Interestingly, there were more variants which were highly correlated (absolute correlation > 0.5) with pseudotime than either of the physiological markers (Figure 5A). To further investigate whether all variants were simply correlated with bolting status, we clustered these variants across pseudotime. We found that there were clusters of variants which transition from one allele to another at different points across pseudotime (Figure 5B). This indicates that genetic variation had a more complicated relationship with the vegetative-to-reproductive transition than a binary determination of ‘bolted’ or ‘non-bolted’. The variants that were most correlated to pseudotime were not co-localised across the genome, indicating that they were not part of the same linkage disequilibrium block. Even though there are only a small number of variants within this near isogenic population, a large proportion of them appear to be correlated with pseudotime, forming 6 clusters of variants, demonstrating that there may be many independent ways of controlling developmental rates. These results further validate the biological relevance of pseudotime, by indicating that genetic variants are more closely associated with it than with directly measured traits, like biomass and leaf size.

**Figure 5:**
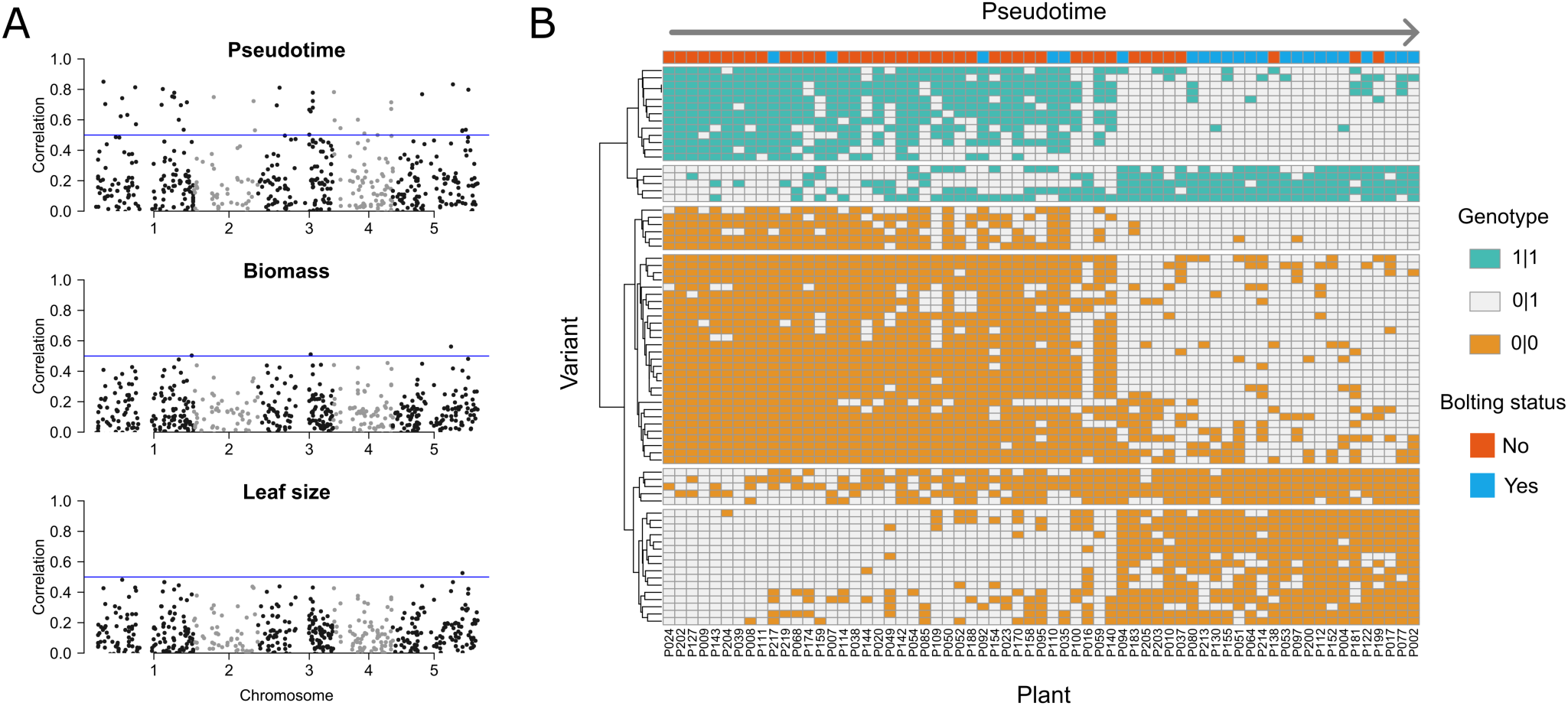
Comparison of variants to pseudotime and physiological traits. Note: in this figure, 0|0 corresponds to homozygous for the reference allele, 0|1 corresponds to heterozygous, and 1|1 corresponds to homozygous for the mutant allele. **A:** Correlation of variants to pseudotime, biomass and leaf size. Correlation was calculated by assigning to 0 to 0|0, 1 to 0|1, and 2 to 1|1, then computing the absolute value of Pearson correlation between genotype and the measurement of interest. The blue line corresponds to a correlation of 0.5. **B:** Genotypes of variants with a high correlation (> 0.5) to pseudotime. Columns (plants) are ordered by increasing pseudotime. Rows (variants) are ordered by hierarchical clustering. To create the splits in the rows, the hierarchical clustering was split into 6 groups.

## Discussion

Single-plant-omics enabled us to identify genes associated with the binary developmental transition and gradual developmental traits. More than half of all Arabidopsis genes significantly change their expression levels at the onset of bolting. By increasing the number of biological replicates, our single plant-omics approach allows us to detect many more differentially expressed transcripts than in previous studies (Woo et al. 2016). In addition, we were able to identify a distinct set of transcriptional regulators that were predictive of leaf size and biomass accumulation. These included several PIFs and circadian clock-related genes, which have been implicated with the regulation of growth rates (Leivar and Monte 2014; Dodd et al. 2005). These results suggest that single-plant-omics can help us distinguish between the transcriptional processes associated with fast and slow developmental processes.

However, even though developmental transitions are rapid, the transcriptional regulation that occurs during these transitions is governed by biochemical processes that will change smoothly over time. One of the main benefits of the single-plant-omics approach is that it enables us to piece together these smooth changes in what initially appears to be a switch-like transition process. We developed a new method of ordering individual plants by their biological age, which we refer to as their pseudotime. We could consistently identify an ordering of plants when utilising independent subsets of genes. In addition, we show that it is biologically meaningful due to the fact that genetic variants in the population of wild type plants are more closely associated with pseudotime than they are to measured traits like biomass and leaf area. While other methods for pseudotime inference are widely used (Saelens et al. 2019; Ding, Sharon, and Bar-Joseph 2022), these are designed for single cell RNA-seq datasets with hundreds or thousands of cells, while our method is effective for smaller sample sizes.

By using pseudotime, we were able to dissect the order of events that took place during bolting at a higher resolution than other studies that relied on RNA-seq time series (Breeze et al. 2011; Woo et al. 2016). Although it is well known that the timing of bolting is closely associated with leaf senescence in different *Arabidopsis* accessions and mutants (Balazadeh et al. 2008; Kim, Park, et al. 2018; Upadhyay et al. 2014; Yan et al. 2017), here we show that many senescence-related processes are concurrent with bolting. Indeed, some of the early processes associated with senescence, such as shutting down ribosome production, appear at a transcriptional level before the visible onset of bolting, which challenges preconceived notions about the order of events that occur during bolting. We identified 43 key transcriptional regulators and identified the order in which they increase or decrease their expression during the vegetative-to-reproductive transition. Our resulting gene network will be a valuable resource for exploring the regulation underpinning this fundamental developmental transition.

Our work suggests that single plant-omics provides numerous benefits for the understanding of rapid developmental transitions, such as (i) increased sensitivity to differentially expressed genes (ii) ability to distinguish between gene expression changes associated with fast and slow developmental processes (iii) capacity to reconstruct the sequence of transcriptional events that occur during a rapid developmental process. Single-plant-omics coupled with pseudotime inference will be widely applicable to investigate other rapid developmental processes, such as the juvenile-to-vegetative transition, the onset of flowering, and the initiation of the vegetative-to-reproductive transition in the meristem.

## Acknowledgements

This project was undertaken on the Viking Cluster, which is a high performance compute facility provided by the University of York. We are grateful for computational support from the University of York High Performance Computing service, Viking and the Research Computing team. We also acknowledge support from the Horticulture Facility and the Technology Facility at the Department of Biology, University of York.

## Funding

This work was supported by funding from the UK Biotechnology and Biological Sciences Research Council (BSSRC): DE/SJD: BB/V006665/1, DE: BB/S506795/1, and SJD: BB/N018540/1. We also acknowledge BBSRC White Rose DTP studentships (BB/M011151/1 and BB/T007222/1) to JR (ref 1792522) and EJR (ref 2444228). DE was also supported by the Royal Society (RGS\R2\212345).

### Author contributions

EJR: Conceptualization; Methodology; Software; Formal analysis; Investigation; Writing - Original Draft; Visualization

JR: Conceptualization; Investigation; Writing - Review & Editing

SJD: Writing - Review & Editing; Supervision; Funding acquisition

DE: Conceptualization; Methodology; Writing - Original Draft; Writing - Review & Editing; Supervision; Funding acquisition

### Data availability

Raw and processed sequencing data have been deposited in the NCBI Gene Expression Omnibus (GEO) database, under accession number GSE242681 (https://www.ncbi.nlm.nih.gov/geo/query/acc.cgi?acc=GSE242681).

Scripts used to produce the figures have been deposited in a GitHub repository (https://github.com/stressedplants/SinglePlantOmics).

## Methods

### Plant growth conditions, library preparation, and RNA-seq

Seeds from the Arabidopsis ecotype Wassilewskija Ws-2 were surface sterilised and plated onto 1x MS plates (1.5% agar) supplemented with 1% w/v sucrose. After four days of stratification at 4°C plates were transferred to long-day photoperiods (16h light/8h dark, 21°C) for 10 days. After 10 days, 225 seedlings were individually transferred to soil (5% sand) and grown for a further 10 days. The 3^rd^ and 4^th^ leaf was tracked during growth. On day 20, 75 plants were selected from the population to ensure a diversity of bolting statuses. For each of these 75 plants, they were classified as bolting if there was a visible bolt ∼1 cm above the rosette, and not bolting otherwise. Images of the 3^rd^ and 4^th^ leaf were then taken for later analysis. On day 21 at ZT4 (4-hours after lights on), the 3^rd^ and 4^th^ leaf were harvested and snap-frozen in liquid nitrogen. The rosette biomass minus the 3^rd^ and 4^th^ leaf was then measured.

Total RNA was isolated from leaf tissue using the Qiagen RNeasy Plant Mini Kit (Cat no. 74904). Residual genomic DNA was removed using the Invitrogen Turbo DNA-free kit (Cat no. AM1907), according to the manufacturer’s protocol. Libraries were prepared with the NEBNext Ultra II Directional Library Prep Kit for Illumina (Cat no. E7765), using the NEBNext poly(A) magnetic isolation module (Cat no. E7490). Quality control was performed with the Agilent 2100 Bioanalyzer instrument (Part no. G2939BA). Finally, a total of 70 libraries were pooled and sequenced, via Novagene, using one lane on an Illumina NovaSeq system. Before analysis of the raw sequencing data, FastQC v0.11.7(Andrews et al. 2012) was used to assess read quality. Illumina adapters were trimmed using CutAdapt v3.4(Martin 2011). Reads were quantified using Salmon v1.6.0 (Patro et al. 2017) and the TAIR10 transcriptome (Berardini et al. 2015).

For further analysis except differential gene expression, transcripts per million (TPM) was used as the measure of relative gene expression across samples. Additionally, TPM levels per transcript isoform were combined to leave only gene-level expression data. One sample (P158) was sequenced again due to low read depth in the initial sequencing and the initial data was discarded. Several samples (P128, P196, P098, P169, P079) were discarded as outliers during initial clustering of the data (Supplementary Figure 1). Finally, TPM values were filtered to remove genes with very low expression, to avoid biassing clustering and other analyses. Genes were removed for low expression if they met either of these two criteria: *(i)* TPM of 0 in 10 or more samples, or *(ii)* TPM below 0.5 in all samples. This left 19208 genes for further analysis. After this filtering, samples clustered into two large groups (Supplementary Figure 2).

### Differential gene expression and GO term overrepresentation

Due to the large number of samples, which may lead to a poor fit to the assumed distributions of parametric tests, we used the non-parametric Mann-Whitney U test to perform differential gene expression analysis (Li et al. 2022). Counts per million (CPM) were used as inputs to this test and p-values were corrected according to the Benjamin and Hochberg method, as suggested by ref. (Li et al. 2022). Genes were classified as differentially expressed if both of the following conditions were true: (i) the adjusted p-value was less than 0.05, and (ii) the log2 fold change between the means with the non-bolted and bolted plants was greater than 0.1 or less than -0.1 (Supplementary Figure 3 and Table S1). GO term overrepresentation was performed using the gprofiler2 R package (version 0.2.1) (Kolberg et al. 2020) (Table S2).

### Physiology trait prediction with Elastic Nets

Due to the large number of predictors (i.e., genes) versus samples, the Elastic Net was chosen to regularise linear models used in prediction, because it incorporates variable selection but also balances for accuracy by allowing more coefficients to be non-zero (Zou and Hastie 2005). TPM values were z-scored before input. A set of known and potential regulatory genes was curated by selecting genes annotated with the following GO terms: “DNA binding transcription factor activity” (GO:0003700), “signal transducer activity” (GO:0004871), and “regulation of transcription - DNA-templated” (GO:0006355), which were the same as those in a previous single-plant-omics data (Cruz et al. 2020). These were used as the set of possible predictors for all figures in the main text.

A tiered cross-validation approach was used to find optimal hyperparameters. This was implemented using ‘scikit-learn‘ in Python v3.10.4 using the ‘ElasticNetCV’ function (Pedregosa et al. 2011). Firstly, leave-one-out-cross-validation (LOOCV) was used to produce a 65 separate test-train splits, where the testing set only consisted of a single sample. Then for each training set, 5-fold cross validation was used to pick optimal ‘alphà and ‘l1_ratiò, with possible inputs for ‘l1_ratiò of 0.1, 0.5, 0.7, 0.9, 0.95, 0.99, and 1.0 (Table S3). The output of this process was a set of optimal hyperparameters and a trained Elastic Net model for each test-train splits. The exact coefficients in the models are summarised in Table S4.

### Pseudotime inference

Since our sample size was too small to infer an ordering of samples using unsupervised techniques, which is typical in single-cell RNA-seq data analysis (Saelens et al. 2019), we developed a new pseudotime inference method. We applied the following steps to genes identified as differentially expressed (see above). First, we processed the gene expression values for each gene as follows: (i) z-score TPM values per gene and truncate any values greater than 3 or less than -3; (ii) linearly scale these values, so that the minimum value was 0 and maximum value was 1 for each gene; and (iii) if the gene was identified as down-regulated, change the gene expression to be 1 - x. To validate this assumption, we ‘bootstrapped’ our pseudotime inference, by partitioning genes into 100 distinct groups and repeating the process for each group. Specifically, define 𝑥_*i,j,k*_ as the normalised expression of the 𝑖th gene in group 𝑗 in the 𝑘th sample, where 𝑗 = 1, …, 100, 𝑖 = 1, …, 𝑛_#_, and 𝑘 = 1, …, 65. Then for each group 𝑗, the 𝑘 samples were ordered based on ∑^*n_j_*^_*i*=1_ *x_i,j,k_*, i.e. the sum of normalised expression of every gene within a fixed group and sample.

There was a consistent pseudotime predicted by each group (Supplementary Figure 5). Finally, we combined the individual predicted pseudotemporal orderings, by assigning each sample to its most common predicted position between the separate orderings. See Supplementary Figure 6 for a visualisation of these results using an unsupervised dimensionality reduction method (principal component analysis; PCA).

Following pseudotime inference, gene expression values were smoothed and filtered based on the goodness of fit of the smoothed curves. Cubic B-splines were used as basis functions to fit to the data and the smoothing process was penalised with the second derivative. The fitting process was repeated, varying the number of basis functions and the smoothing parameter (‘lambd’), to choose optimal hyperparameters, i.e. those that minimised the sum of the generalised cross-validation (GCV) across all genes (Kokoszka and Reimherr 2021) (Supplementary Figure 7A; Table S5). Finally, the 4000 genes with lowest GCV were chosen for further analysis (Supplementary Figure 7B; Table S6). The smoothed functions were again rescaled to ensure they had a minimum of 0 and maximum of 1, allowing for comparisons between timeseries and tolerance to outliers. Additionally, some genes had a few samples where the TPM was much higher than in other samples and these genes had inappropriately smoothed curves. Genes were therefore removed if the ratio between their range and their interquartile range was too high (Supplementary Figures 7C and 7D; Table S6). 3928 genes remained after these filtering steps.

### Analysis of gene expression over pseudotime

We summarised the expression dynamics of each monotonically increasing or decreasing gene by its area under the curve (AUC). Firstly, we filtered for genes whose smoothed expression was monotonic over pseudotime. Specifically, a gene was considered monotonic if the derivative of its B-spline-fitted expression over pseudotime (see “Pseudotime Inference”) never crossed 0. (Due to the potential for numerical errors when calculating the derivative, we set all derivative values within +/-1e-3 to 0, to avoid spuriously removing monotonic genes.) Then, we summarised these genes by calculating their AUC - i.e. the integral of the curve between the start and end of pseudotime. Crucially, since these selected genes are monotonic, a decreasing gene with smaller AUC changes more rapidly earlier on in pseudotime (see Figure 3C). For increasing genes, the AUC was calculated for (1 - normalised gene expression) values.

After we calculated these values, we chose to group genes by GO term, to find which GO terms had significantly different average AUC values. (See Supplementary Figure 8 and Table S7 for details about selecting thresholds for GO term size, used before making comparisons.) We used the non-parametric Kruskal-Wallis test to see if there were any significant differences between GO terms, i.e. if the p-value was smaller than 0.01. This was then followed by a non-parametric post-hoc test to find significantly contrasting GO terms, i.e. where the corrected p-value was smaller than 0.01. Specifically, we used the command ‘kwAllPairsNemenyiTest’, with the Chi squared approximation, from the PMCMRplus v1.9.7 R package (Pohlert 2023; Nemenyi 1963). Adjusted P-values (via the ‘single-step’ method) are summarised for every test in Tables S8 and S9. Additionally, we visualised the distribution of AUC values for all GO terms within the same size boundaries (Supplementary Figures 9 and 10).

### Gene regulatory network analysis

After pseudotime inference, we applied a gene regulatory inference method (DynGENIE3) designed for timeseries gene expression data (Huynh-Thu and Geurts 2018). Since this method required a prespecified list of potential regulatory genes, we selected TFs included in a previous DAP-seq experiment (O’Malley et al. 2016). In addition, we filtered for TFs which passed pseudotime filtering (see “Pseudotime inference”). This resulted in a potential 46 transcription factors. After running DynGENIE3, we kept only edges with a predicted weight of at least 1e-10, then further selected the top 5% of all edges by weight (Supplementary Figure 11).

To select for high-quality predicted interactions, we filtered the results against the DAP-seq dataset (O’Malley et al. 2016). Specifically, the putative regulatory targets (with FRiP greater than or equal to 5%) were downloaded for all sequenced TF binding sites. This was retrieved from http://neomorph.salk.edu/dap_data_v4/fullset/dap_download_may2016_genes.zip on 01/06/2023, via http://neomorph.salk.edu/dap_web/pages/browse_table_aj.php. Overall, this GRN included 1668 genes and 2248 interactions (Table S10).

For graph visualisation, Gephi v0.9.7 was used (Bastian, Heymann, and Jacomy 2009). Note that for Figure 4, not all TFs were monotonic over pseudotime (see “Analysis of gene expression over pseudotime”). Therefore, a TF was classified as ‘decreasing’ if the initial value of the smooth curve over pseudotime was greater than the final value and classified as ‘increasing’ otherwise.

### Variant calling

To analyse genetic variation within the population, we followed GATK guidelines for short variant discovery from RNA-seq data. We used STAR (v2.7.10b; two-pass mode) to align reads to TAIR10 genome, revision 56 (Dobin et al. 2013). We then pre-processed the aligned reads using the commands ‘MarkDuplicates’ and ‘SplitNCigarReads’ from GATK (v4.3.0.0) (Poplin et al. 2018). We used the command ‘HaplotypeCaller’ to produce genomic variant calling format (gVCF) files per sample. Finally, we produced identified single-nucleotide variants (SNVs) and insertions / deletions (indels) which were confidently called across the whole population, using ‘GenotypeGVCFs’.

For further processing of these initial SNVs and indels, we followed the filtering guidelines suggested by ref. (Cruz et al. 2020). Specifically, we selected only biallelic variants with a minimum genotype quality of 40, and which were called in at least 80% of all samples, using VCFtools (v0.1.16) (Danecek et al. 2011). We then imputed missing genotypes using Beagle (v5.4, 22Jul22, 46e) on default settings (Browning, Zhou, and Browning 2018). Finally, we selected variants with a minor allele frequency (MAF) of at least 0.05.

Note, due to pre-processing by Beagle, the variant calling format (VCF) file includes two versions of the same heterozygous haplotype (‘0|1’ and ‘1|0’) (Table S11; (Browning, Zhou, and Browning 2018)). In processing for Figure 5, these were combined into a single heterozygous haplotype, and any variants with only heterozygous haplotypes across all samples were removed.

## Supplementary Materials

### Description of predictors identified in Elastic Net analysis

We were able to identify genes that were consistently identified as predictors of both leaf area and biomass in multiple models during the LOOCV process. Overall, 702 regulatory genes had a non-zero median coefficient across all models (Figure 2C). Of these, 59 were identified as positive predictors of biomass and leaf size and 86 were identified as negative predictors of both traits (Figure 2C). The gene with the highest positive coefficient for both traits was HOMEOBOX PROTEIN 54 (HB54; AT1G27045), involved in a nitrogen-signalling cascade linked to plant growth (Ariga et al., 2022). Two NAC (no apical meristem, NAM; Arabidopsis transcription activation factor, ATAF; cup-shaped cotyledon, CUC) family transcription factors had high positive coefficients for both traits: XYLEM DIFFERENTIATION, DISRUPTION OF VASCULAR PATTERNING (XVP; AT1G02220) is involved in vascular development (Yang et al., 2020); and NAC4 (AT5G07680), is involved in pathogen-induced cell death (M.-H. Lee et al., 2017). Similarly, we identified genes with highly negative coefficients for both traits. TARGET OF EARLY ACTIVATION TAGGED (EAT) 2 (TOE2; AT5G60120) is a highly negative predictor of both traits, suggesting that there is a link between leaf size, biomass, and the microRNA/SPL-mediated juvenile to mature transition (Werner et al., 2021). Other highly negative predictors of both traits include transcription factors controlling leaf wax biosynthesis (MYB-SHAQKYF 2; MYS2; AT2G40260; (Liu et al., 2022)) and photomorphogenesis (PHYTOCHROME-INTERACTING FACTOR 7; PIF7; AT5G61270; (Burko et al., 2022)).

We next sought to understand important predictors of one trait which were not predictive of the other trait. Interestingly, a pair of genes involved in nutrient transport within phloem (Wei et al., 2021), NAM (AT1G52880) and NAC56 (AT3G15510), were highly positively predictive of biomass but not of leaf size. WRKY2 (AT5G56270), a regulator of PHYTOCHROME-INTERACTING FACTOR 4 (PIF4; AT2G43010; (Wang et al., 2022)), was also a positive predictor of biomass but not leaf size. Conversely, two of the most positive predictors of leaf size which were not predictive of biomass were CONSTANS-LIKE 3 (COL3; a regulator of photomorphogenesis and flowering time; AT2G24790; (Tripathi et al., 2017)) and REVEILLE 7 (RVE7; a circadian-clock- and temperature-responsive gene; AT1G18330; (Tian et al., 2022)). Another central circadian-clock gene, PSEUDO-RESPONSE REGULATOR 3 (PRR3; AT5G60100; (Para et al., 2007)), which is also highly expressed in vasculature, was a strong negative regulator of leaf size but not predictive of biomass. Two leaf-specific transcription factors were strong negative regulators of leaf size but not predictive of biomass: MYB-SHAQKYF 1 (MYS1; involved in leaf wax biosynthesis; AT2G38300; (Liu et al., 2022)) and MYB88 (involved in stomatal development; AT2G02820; (E. Lee et al., 2013)). It is worth noting that comparatively few regulatory genes had an opposite-sign effect on the two traits (5 for positive-biomass/negative-leaf and 8 for positive-leaf/negative-biomass). Of those, a key transcription factor in vernalisation, VERNALIZATION2 (VRN2; AT4G16845; Figure 2D; (Gendall et al., 2001)), was a slightly positive biomass predictor and a slightly negative leaf size predictor.

### Summary of Supplementary Tables

Table S1 - Differentially expressed genes

This contains all the genes identified as differentially expressed (see “Differential expression and GO term overrepresentation” in main text). The columns are:

- id = the TAIR code for the gene
- log2FoldChange = the log2-transformed fold change (i.e. mean of gene expression within bolted plant divided by the mean within non-bolted plants)
- p_adj = the adjusted P value for the Wilcox text between non-bolted and bolted plants
- diffexpressed = “up” if the gene is up-regulated and “down” if the gene is down-regulated
- gene_name = the name of the gene as found by biomaRt

Table S2 - gprofiler results

Results from gProfiler to find overrepresented GO terms within UP and DOWN groups.

Table S3 - hyperparameters

We used a hierarchical cross-validation approach to validate our models to predict physiological data from gene expression data (see “Physiology trait prediction with elastic nets” in Methods for details). This table contains the optimal parameters chosen within the 5-fold cross-validation, after each sample was individually removed from the training data. The different Excel sheets contain these values for the four different Elastic Nets (leaf size or biomass, with all genes or only with regulatory genes).

Table S4 - coefficients in optimal models

Similar to Table S3, these contain the coefficients and summary statistics for each of the optimal models chosen by 5-fold cross validation. Again, different sheets contain these values for the four different Elastic Nets (leaf size or biomass, with all genes or only with regulatory genes).

Table S5 - GCV values with different hyperparameters

This table contains the sum of GCV across all genes, when fitting B-splines with varying numbers of basis functions and applying a different level of regularisation (“Pseudotime inference” in Methods; also visualised in Supplementary Figure 7A). The optimal hyperparameters chosen were: 9 basis functions and a regularisation parameter of 1178.769, since these values produced the lowest sum of GCV across all genes.

Table S6 - values for filtering out genes, based on smoothing

This table contains values which were used to filter for genes which varied smoothly over pseudotime (“Pseudotime inference” in Methods). ‘gcv_value’ corresponds to the GCV of the B-spline fit to each gene (visualised in Supplementary Figure 7B). ‘iqr_value’ corresponds to the ratio between the range and interquartile range of the scaled TPM data (not the smoothed values; visualised in Supplementary Figure 7C).

Table S7 - Manually defined thresholds for the sizes of GO terms

This table specifies the boundaries between GO terms containing medium and high numbers of genes. The GO term category (molecular function, biological process, cellular component) is specified and whether decreasing or increasing genes were selected. The same boundaries, overlaid with the sizes of individual GO terms, are presented in Supplementary Figure 8.

Table S8 - Statistics for decreasing GO terms

Individual Excel sheets contain the p-values was two types of statistical tests, which were applied to GO terms within GO term categories and of similar sizes (see “Analysis of gene expression over pseudotime” in Methods). “kw_p_value” represents the result for the Kruskall-Wallis test. For the values of the “kwAllPairsNemenyiTest” test, each adjusted p-value is reported between each pair of compared GO terms (Pohlert 2023; Nemenyi 1963).

Table S9 - Statistics for increasing GO terms

This has the same structure as Table S8, but contains information about increasing GO terms.

Table S10 - Link list for GRN

This table contains the list of edges and related edge weights for the final GRN. The columns include:

- ‘weight’ for the relative weight of the edge, as specified by the DynGENIE3 package (Huynh-Thu and Geurts 2018).
- ‘regulatory gene’ for the source node of the edge. The values will be one of the 46 possible transcription factors.
- ‘target gene’ for the target node of the edge.

Table S11 - Variant calling file

This file is a standard variant calling format (VCF v4.2) file, containing information about the haplotype of each sample, for the final filtered variants.

Table S12 – Processed RNA-seq values

Each Excel sheet contains information about gene expression values, where rows indicate genes labelled by TAIR codes and columns indicate the samples. In summary:

- ‘TPM unfiltered’ is the summary of the predicted transcripts per million (TPM) values from Salmon, after being summed across isoforms per gene (see Plant growth conditions, library preparation, and RNA-seq in Methods).
- ‘TPM filtered’ contains the same values, but only for genes which passed initial filtering (Supplementary Figure 1).
- ‘Normalised – pseudotime’ contains the normalised TPM values for pseudotime inference and the columns are ordered by increasing pseudotime (see Pseudotime inference in Methods).

### Supplementary Figures

**Supplementary Figure 1:**
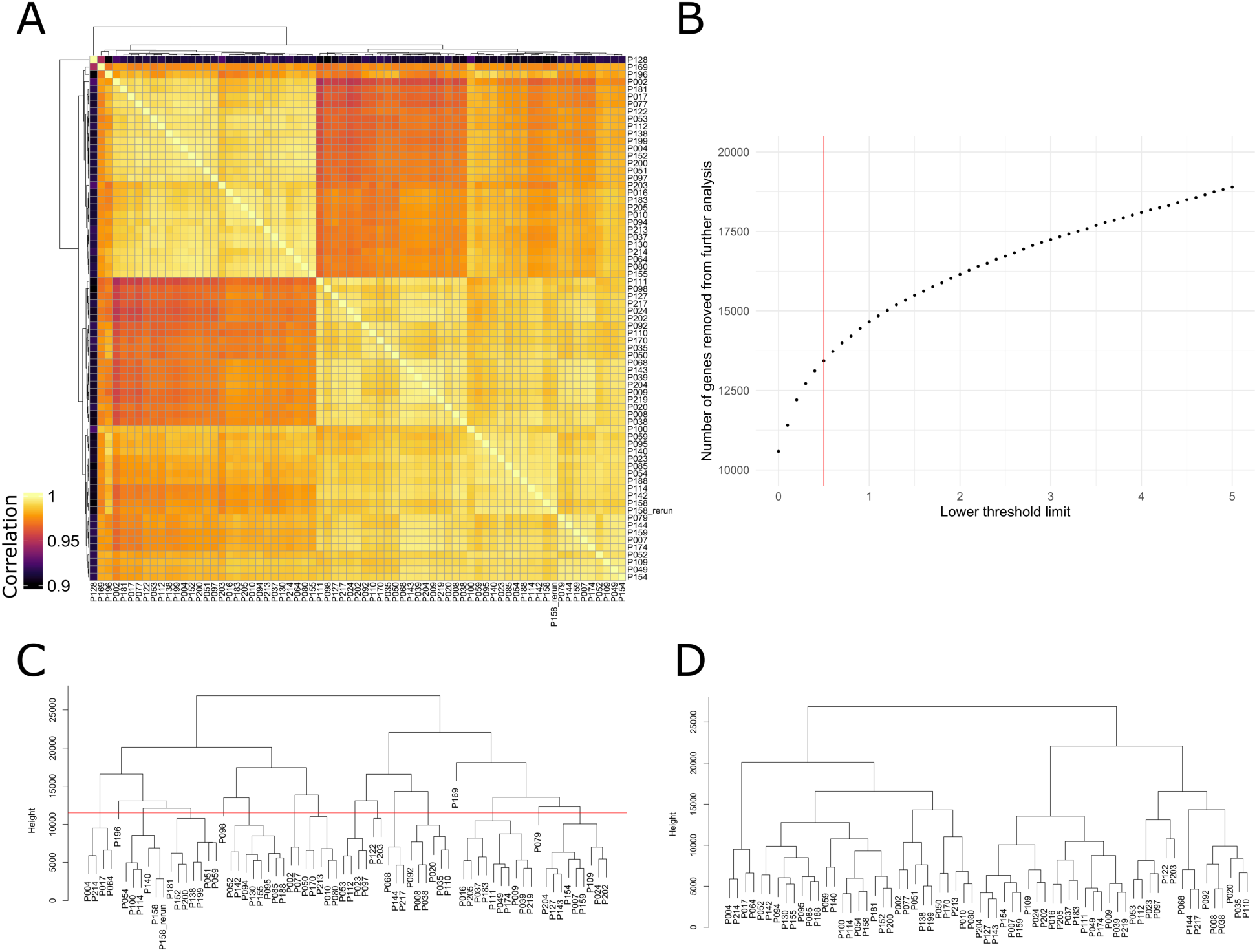
Plots related to filtering of the RNA-seq dataset. **A**: Pearson correlation of each sample based on the entire unfiltered transcriptome (after log transformation). P128 is a clear outlier, with low correlation with every other sample. **B**: The number of genes which would be filtered out based on different low expression thresholds. **C**: Hierarchical clustering (with the complete linkage method) of samples based on TPM values, after the initial removal of sample P128 and low-expressed genes. Four further samples were removed (P196, P098, P169, and P079), since they were separated into single-sample groups by branch splits above a height of 11 500. Additionally, samples P158 and P158_rerun cluster into one group, justifying the replacement of P158 with P158_rerun. **D**: Hierarchical clustering, as in **C**, for the final filtered dataset.

**Supplementary Figure 2:**
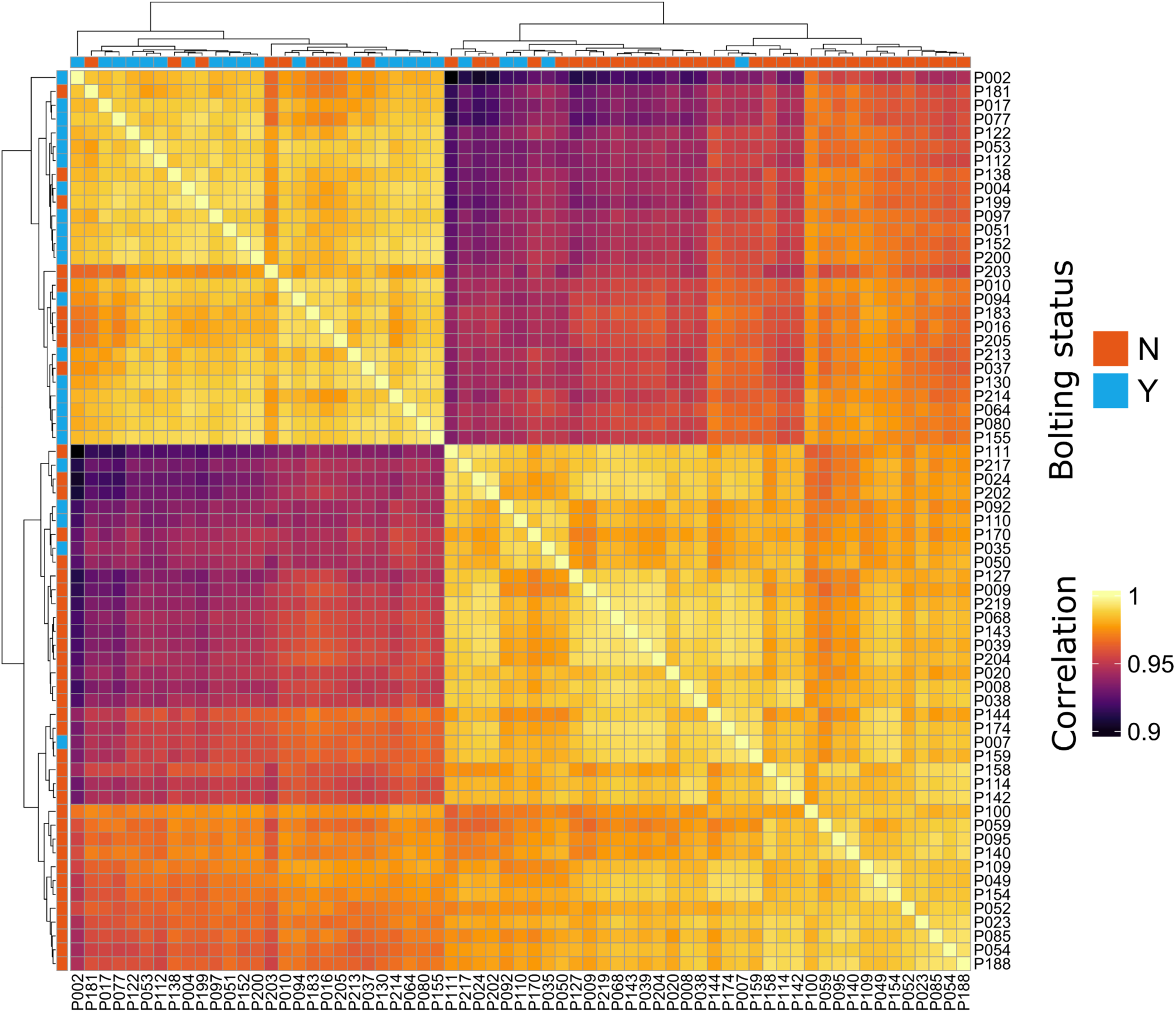
Clustering of samples, based on correlation of log-scaled TPMs. All comparisons between samples show a Pearson correlation coefficient > 0.89. Hierarchical clustering (with the ‘complete’ linkage method) shows that the samples can be separated into two clusters. One cluster contains mostly non-bolted plants and the other contains mostly bolted plants.

**Supplementary Figure 3:**
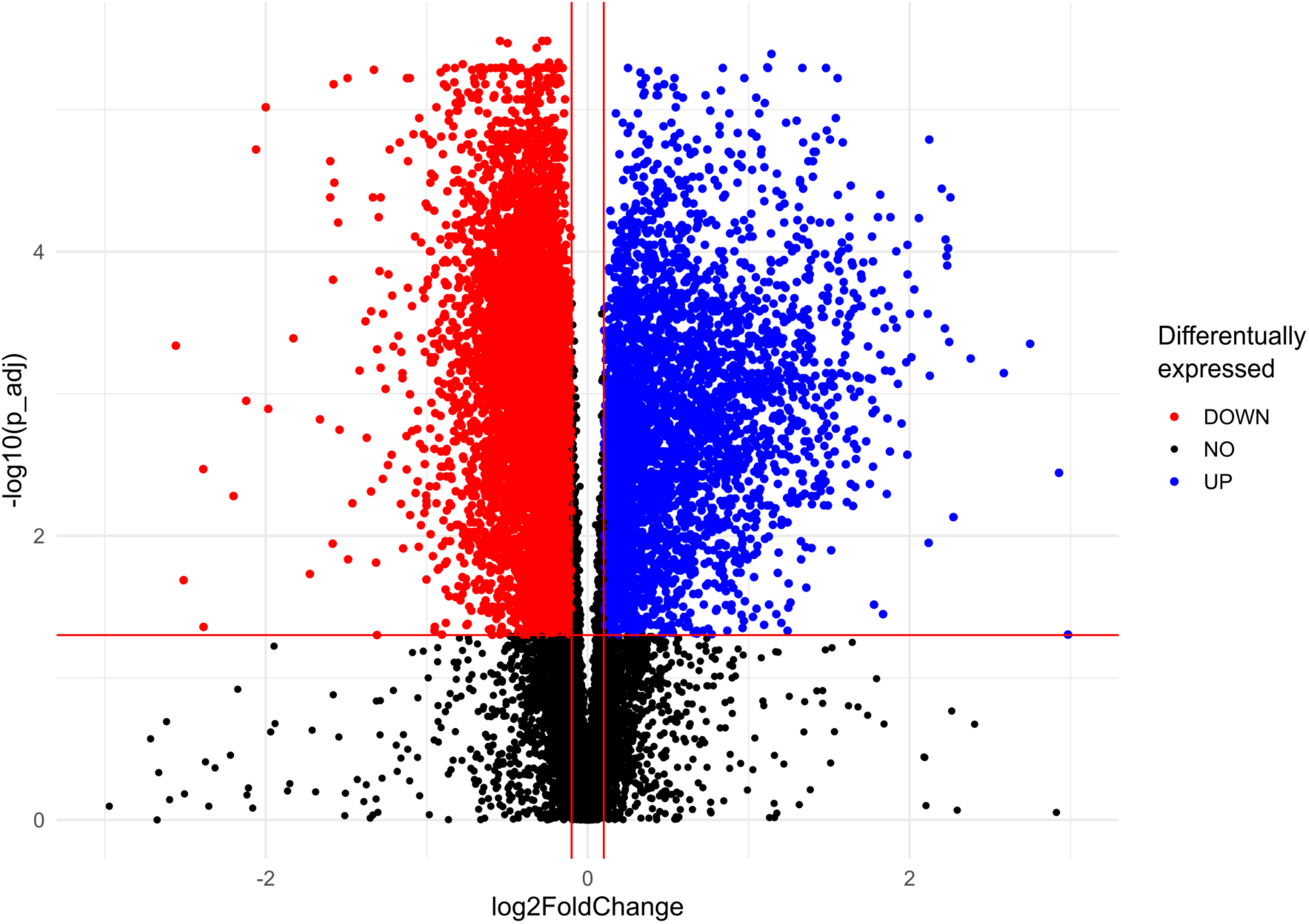
Volcano plot to show differentially expressed genes, based on the bolting status phenotype. We identified genes that: (i) had an adjusted p value of at least 0.05 and (ii) showed a log fold change of more than 0.1 or less than -0.1. (Red lines indicate the boundaries for these decisions.) Out of the 19 208 genes used in this analysis, we identified 6 810 down-regulated genes and 3 754 up-regulated genes (see Table S1).

**Supplementary Figure 4:**
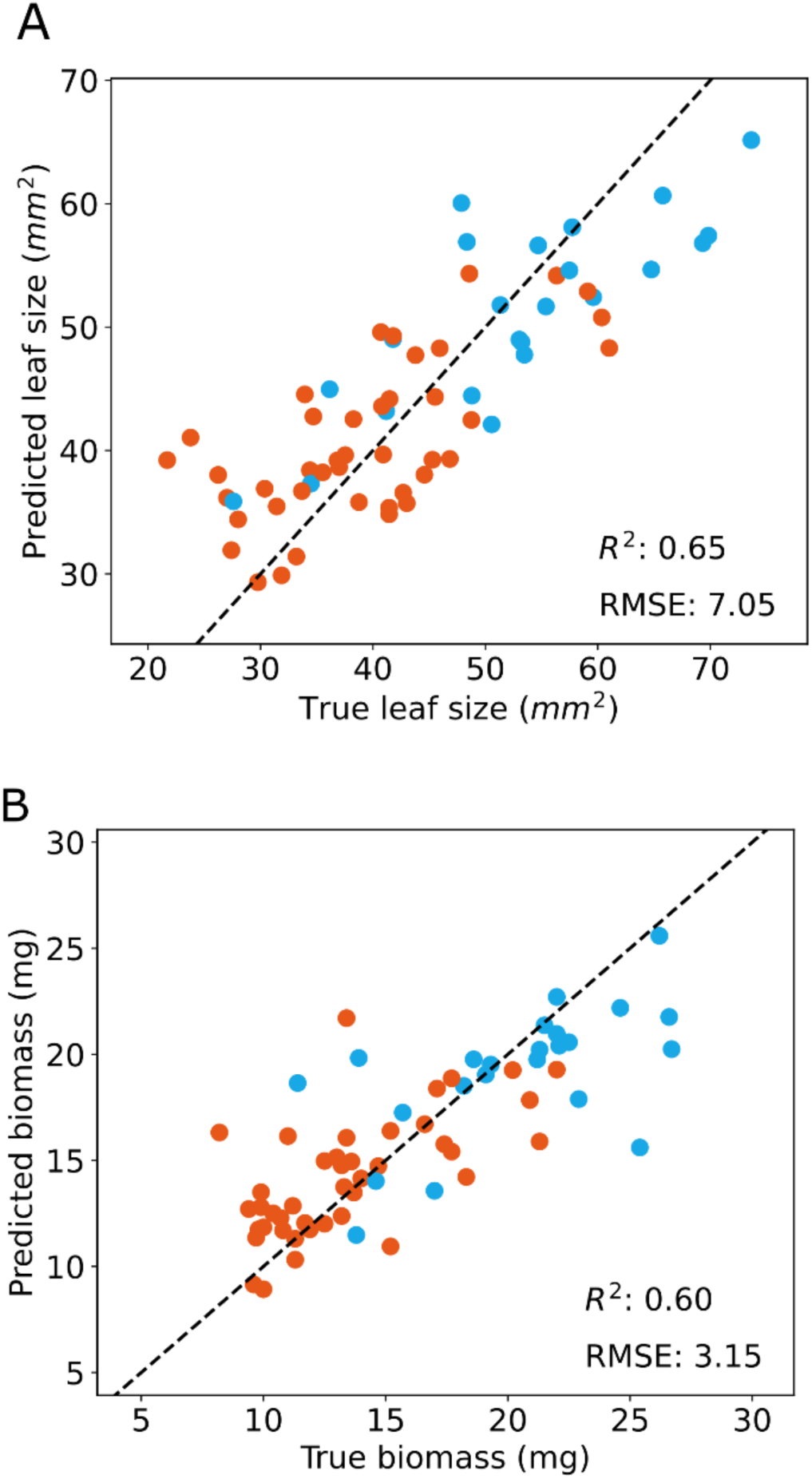
The same as Figures 2A and 2B from the main text, except these models included all genes as potential predictors. For both **A** and **B**, R&#x005E;2 and RMSE metrics are slightly worse than the model with only regulators. Thus, we used the models with a small set of predictors for further analysis in the main text.

**Supplementary Figure 5:**
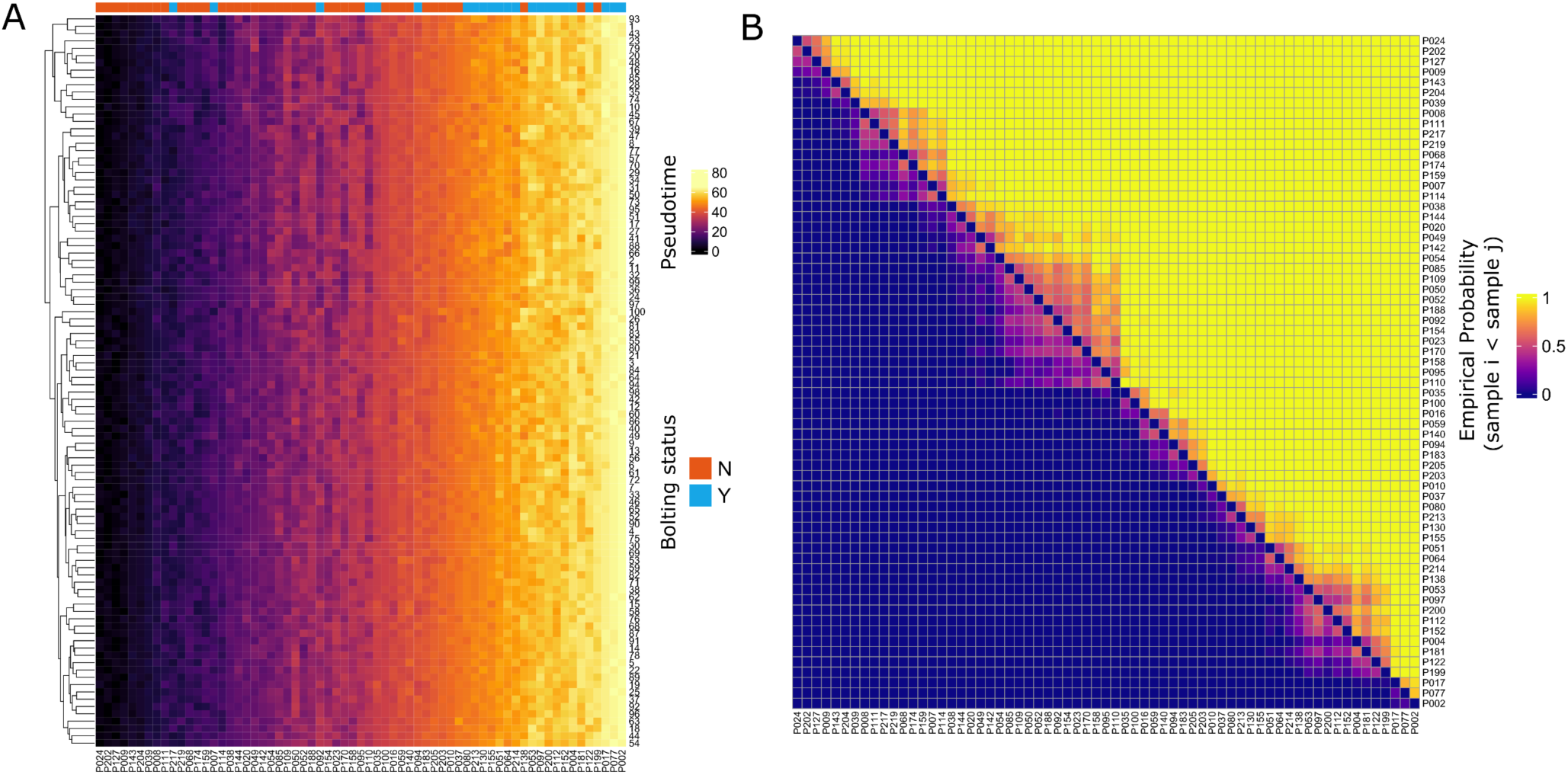
Summary of the bootstrapping process used to infer pseudotime. **A**: Our pseudotime inference method produces consistent results across different groups of input genes for most samples. Values correspond to predicted pseudotime for each sample (columns), as predicted within each partition of the genes (rows). Columns are ordered by the final consensus pseudotime. **B**: There are samples which show some uncertainty in pseudotime prediction - as shown by the clusters of squares around the diagonal. Values correspond to the empirical probability (proportional to the number of observed events in bootstrapping) that the sample of that row is earlier than the sample of that column. Rows (up-to-down) and columns (left-to-right) are ordered by consensus pseudotime.

**Supplementary Figure 6:**
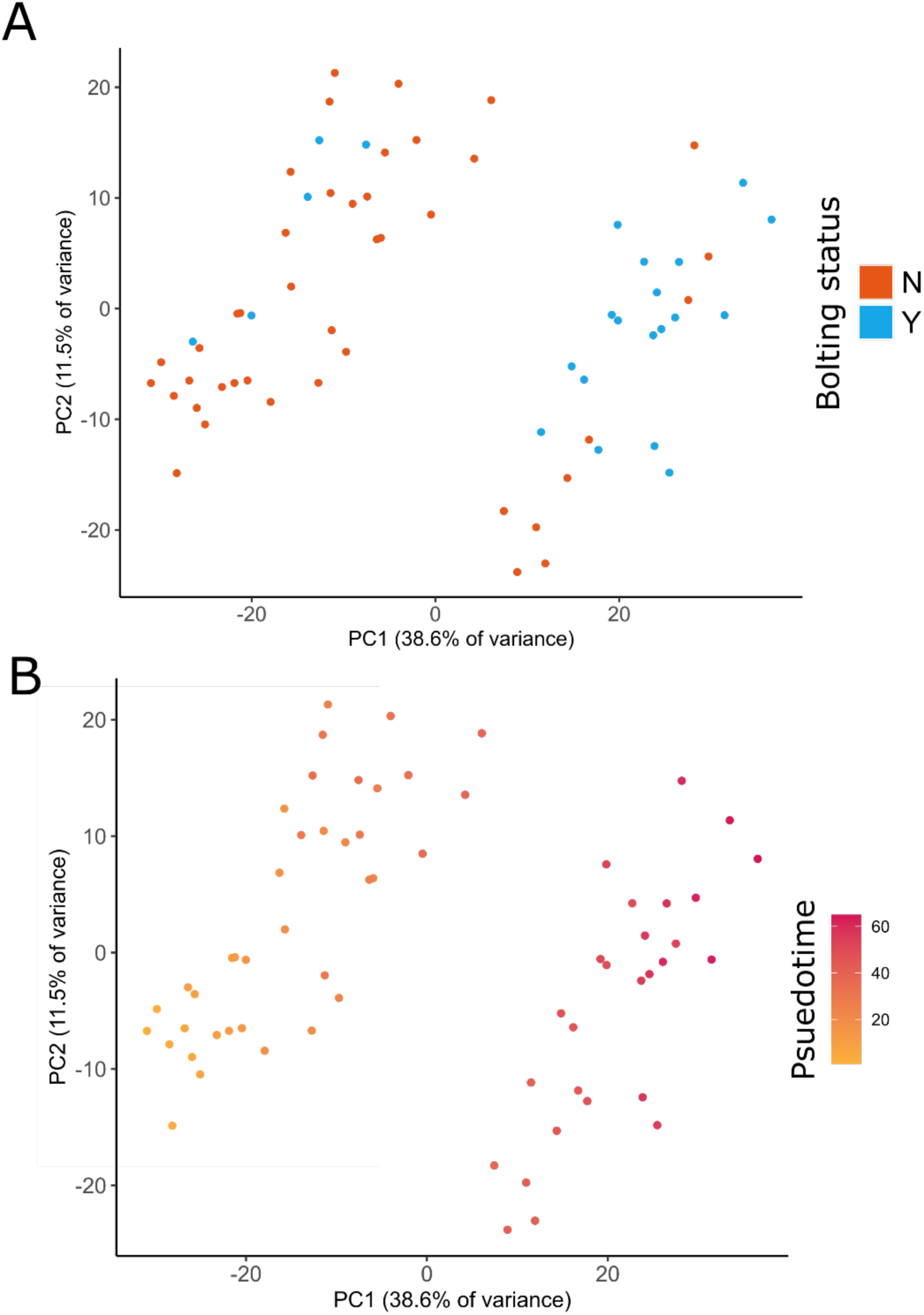
**A**: PC1 separates out the samples into two groups - similar to the groups found in the clustering in Figure 1. **B**: Comparison of our pseudotime inference method with a Principal Component Analysis plot of the gene expression data. *Note*: the PCA was performed after data had been normalized for pseudotime inference.

**Supplementary Figure 7:**
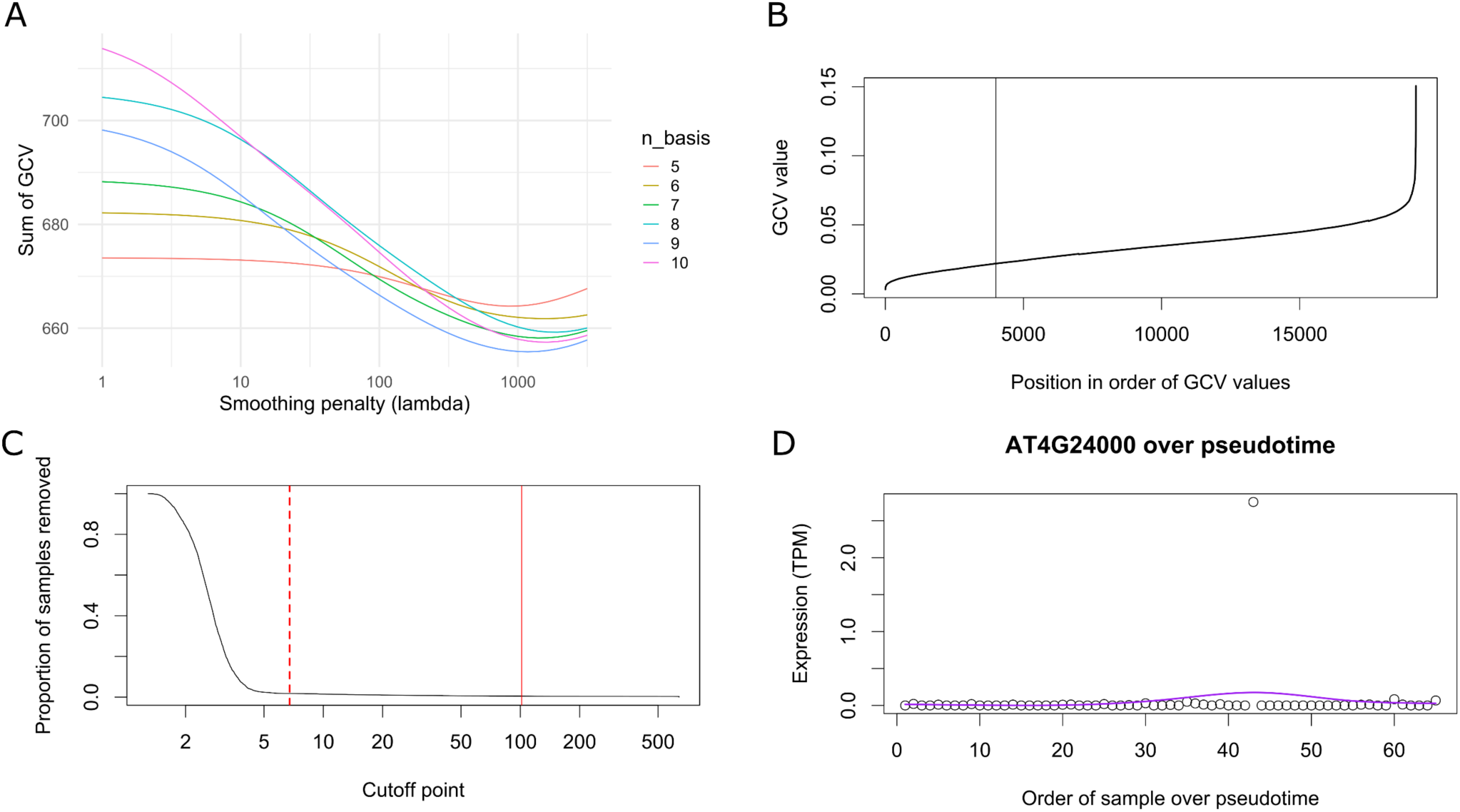
**A**: Selection of optimal parameters for B-spline smoothing. Values correspond to the sum of GCV (generalised cross-validation) across all genes, after B-spline fitting with the fixed number of basis functions (n_basis) and the value of the second derivative smoothing penalty (lambda). *Note:* for reference, the optimal parameters chosen were n_basis = 9 and lambda = 1178.769. **B**: Selection of threshold based on GCV value of each gene. All genes which passed initial filtering are displayed. Genes with high GCV are assumed to not vary over pseudotime and they are therefore removed before further analysis. We selected the 4000 genes with lowest GCV (indicated by the vertical line). **C**: Selection of threshold for interquartile range. Some genes had low GCV, despite displaying poor behaviour over pseudotime (for example, only 1 or 2 samples with high TPM). These were removed by only keeping genes with a low range to interquartile range ratio. Genes with a ratio lower than 6.75 were removed (indicated by the dashed red line). The solid red line corresponds to AT4G24000. **D**: An example of a gene (AT4G24000) removed by the previous filtering. This gene had a ratio of 101.624. The purple line is the B-spline smoothed curve and the points represent TPM values in each sample.

**Supplementary Figure 8:**
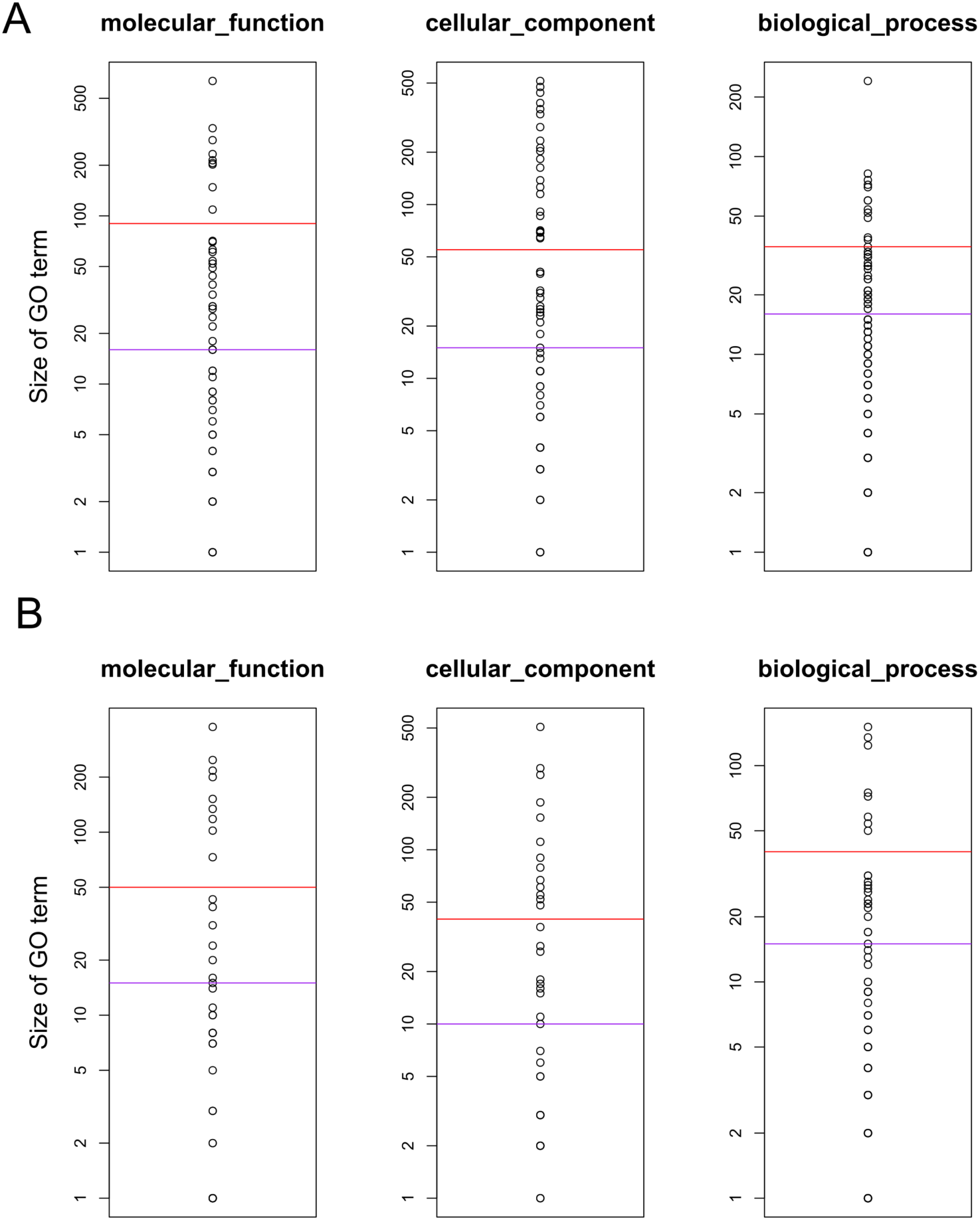
In order to compare ‘area under the curve’ values between groups of GO terms, we first needed to select GO terms of approximately the same size (in terms of the genes which passed all previous filtering steps). Thus, we manually selected thresholds for ‘large’ (between 35 and 90 genes, depending on condition, red line) and ‘medium’ (between 10 and 16 genes, depending on condition, purple line) GO terms. Also, these values needed to be chosen independently for decreasing and increasing genes. **A**: Thresholds chosen for decreasing genes, per GO term. **B**: Thresholds chosen for increasing genes, per GO term

**Supplementary Figure 9:**
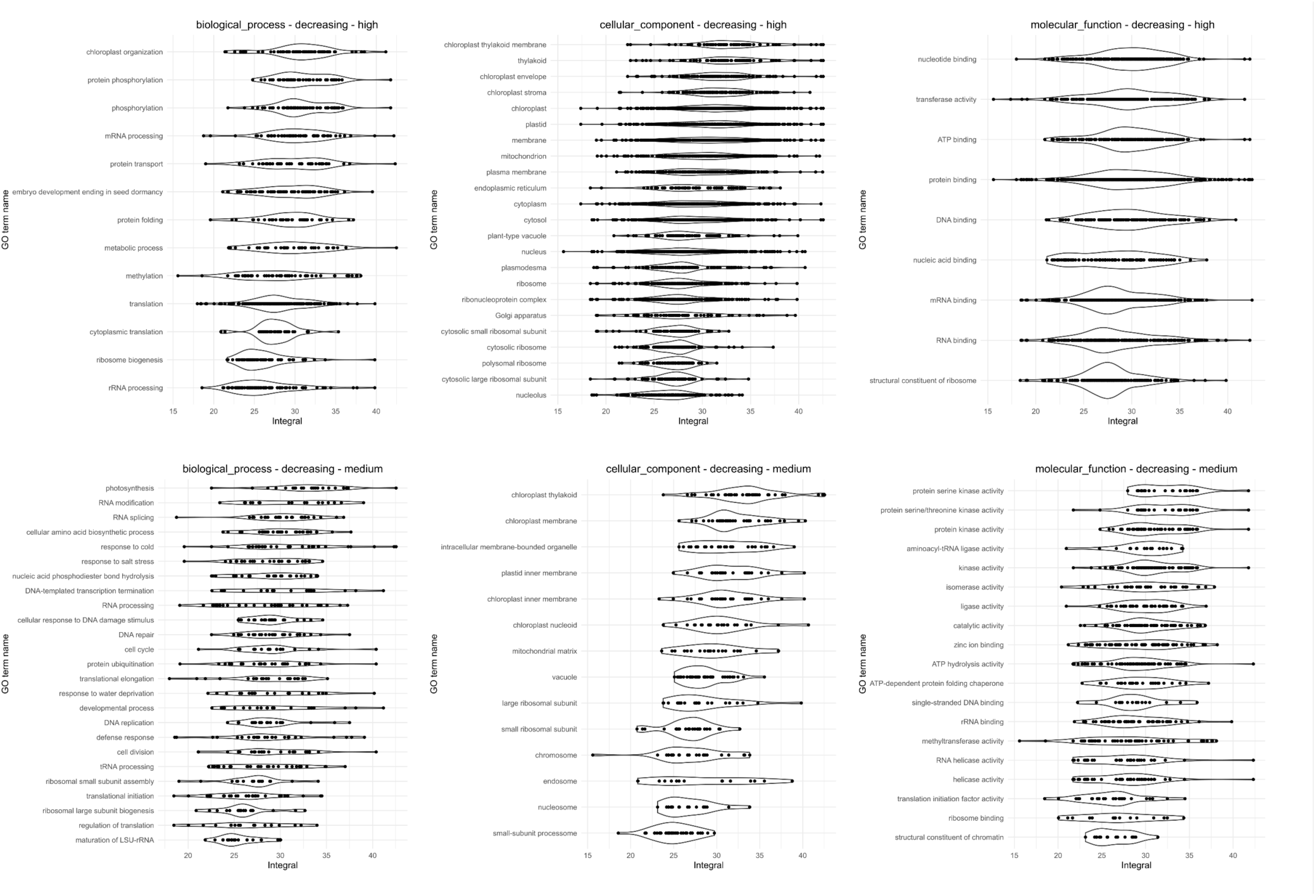
Area under the curve values for GO terms. Filtered to only include genes which decrease over pseudotime. Each violin plot is titled by the category of GO term (molecular function, cellular component, or biological process) and the relative size of the GO term (high or medium; see Supplementary Figure 8). *Note:* Kruskal-Wallis tests and the related post-hoc tests were performed. See Table S8 for details.

**Supplementary Figure 10:**
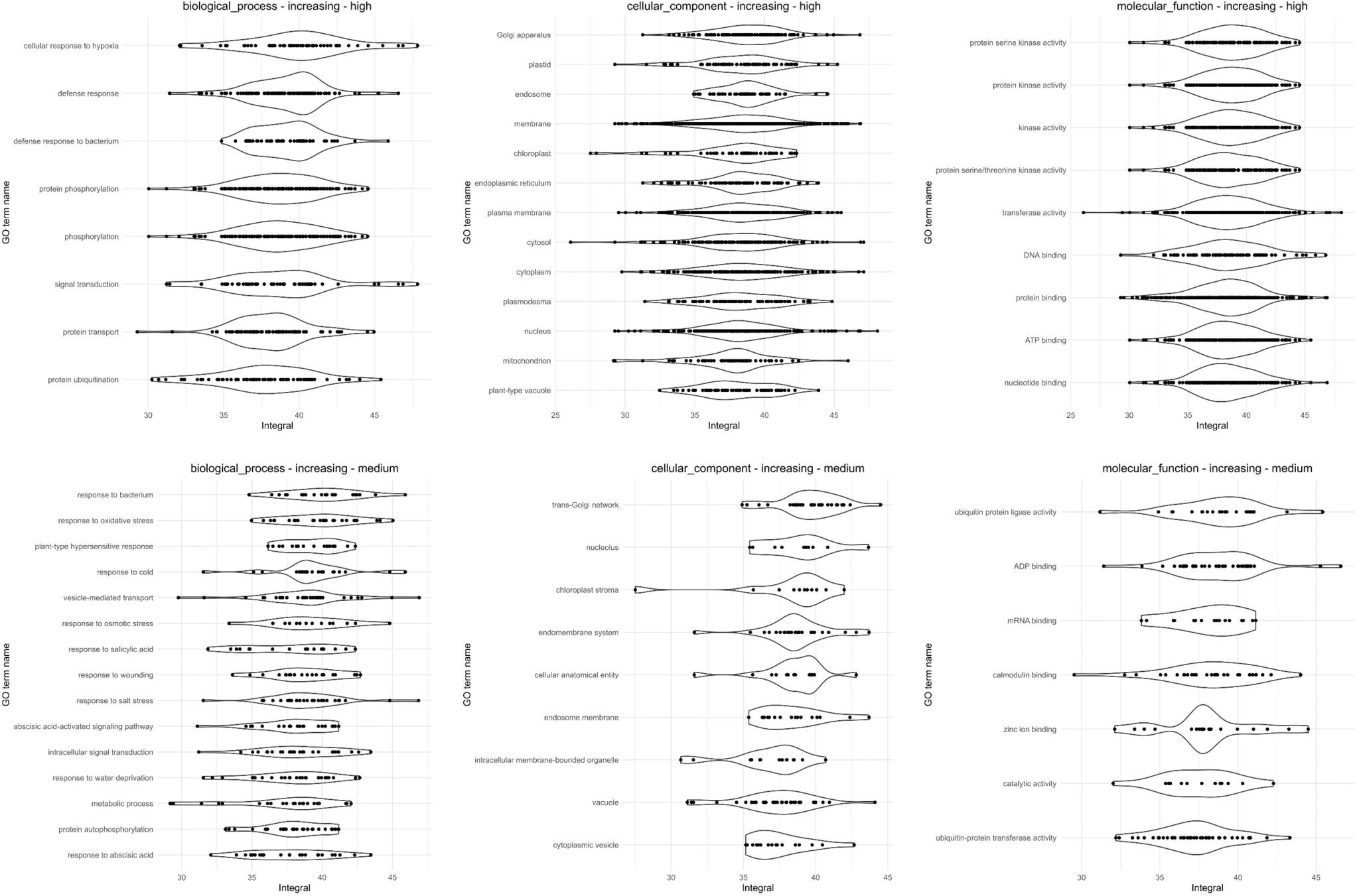
Identical structure to Supplementary Figure 9, except these are for increasing genes. *Note:* most of the Kruskal-Wallis tests and all of the post-hoc tests were not significant (adjusted p value > 0.01) in each of these cases. See Table S9 for details.

**Supplementary Figure 11:**
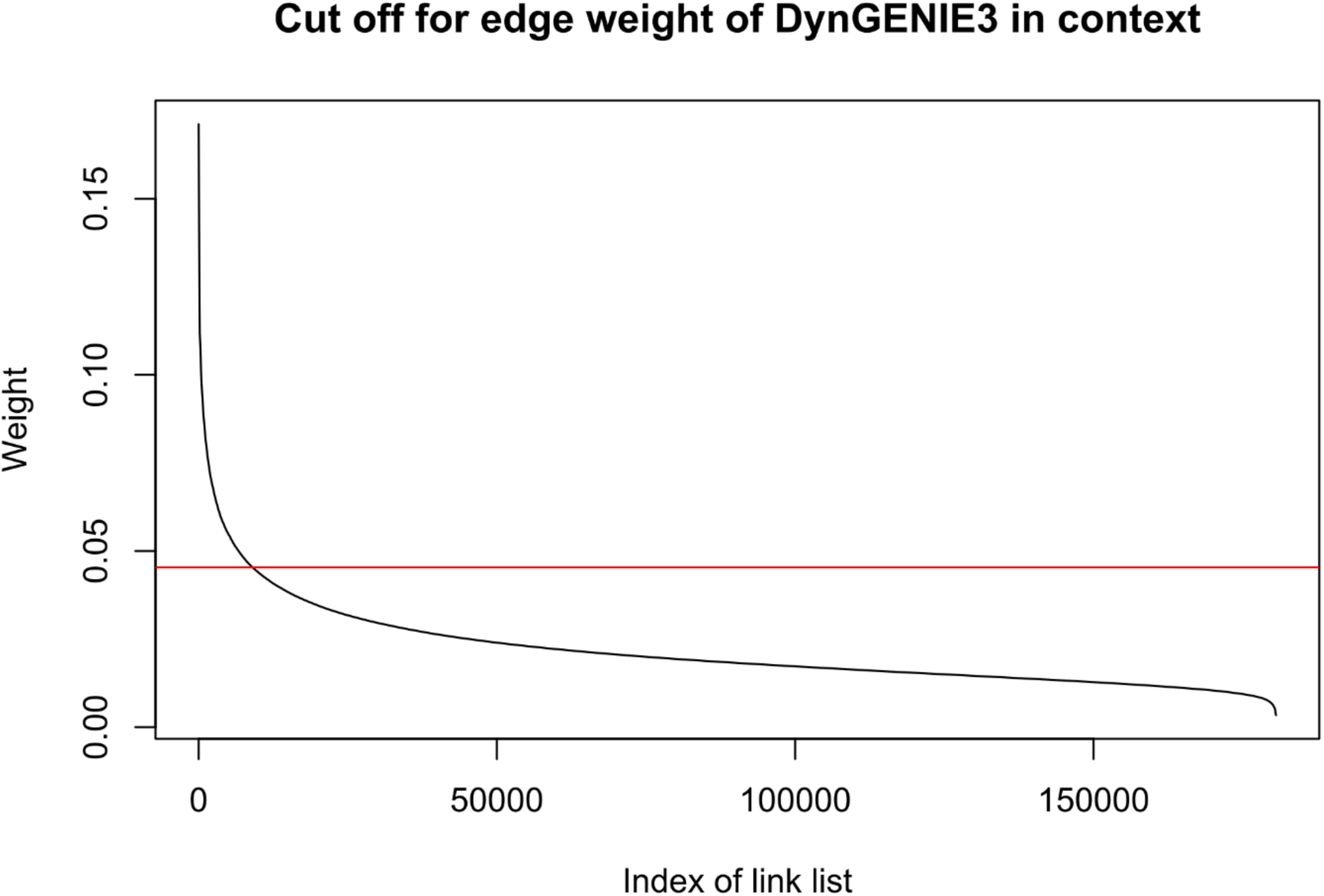
Edge weights of predicted gene-gene interactions by DynGENIE3. The ‘index of the link list’ is defined by decreasing edge weight. The top 5% of edges by weight were included in the final network. *Note:* these weights are only meaningful in a relative sense and have no other statistical significance.

